# A Catalytically Inactive Protein Kinase C alpha Mutation Drives Chordoid Glioma by Pathway Rewiring

**DOI:** 10.1101/2025.05.30.657104

**Authors:** Charlotte Bellamy, Hannah Tovell, Selina Schwaighofer, Timothy R. Baffi, Janan Arslan, Quentin Letourneur, Florent Dingli, Damarys Loew, Alexandr Kornev, Julie Lerond, Tiffany H. Kao, Stephane Liva, Brigitte Izac, Muriel Andrieu, Homa Adle-Biassette, Jean-Vianney Barnier, Eduard Stefan, Marc Sanson, Alexandra C. Newton, Franck Bielle

**Affiliations:** Sorbonne Université, Institut du Cerveau - Paris Brain Institute - ICM, CNRS, Inria, Inserm, équipe labellisée Ligue Nationale contre le Cancer, Hôpital de la Pitié Salpêtrière, F-75013, Paris, France; Department of Pharmacology, University of California, San Diego, La Jolla, CA 92093, U.S.A; Institute of Molecular Biology and CMBI, Technikerstr.25a, University of Innsbruck, Innsbruck, Austria. Tyrolean Cancer Research Institute (TKFI), Innrain 66, 6020, Innsbruck, Austria; Biomedical Sciences Graduate Program, University of California at San Diego, La Jolla, CA 92093, USA; AP-HP.Sorbonne Université, SIRIC Curamus, Paris, France; CurieCoreTech Mass Spectrometry Proteomics, Institut Curie, PSL Research University, 75248 Paris, France; Inserm, U900, Institute Curie, 75005 Paris, France; CNRS, INSERM, Institut Cochin, Université Paris Cité, Paris, France; AP-HP, Hôpital Lariboisière, Department of Pathology, Paris, France; Université Paris-Saclay, CNRS, Institut des Neurosciences Paris-Saclay, 91190, Gif-sur-Yvette, France; Sorbonne Université, Inserm, CNRS, UMR S 1127, Paris Brain Institute - Institut du Cerveau (ICM), équipe labellisée Ligue Nationale contre le Cancer, AP-HP, Hôpitaux Universitaires La Pitié Salpêtrière - Charles Foix, Service de Neurologie 2-Mazarin, Paris, France; Sorbonne Université, Inserm, CNRS, UMR S 1127, Paris Brain Institute - Institut du Cerveau (ICM), AP-HP, Hôpitaux Universitaires La Pitié Salpêtrière - Charles Foix, Laboratoire de Neuropathologie Raymond Escourolle, Paris, France

**Author notes:** Corresponding Author: Franck Bielle, Alexandra Newton. Pfizer Inc, 10555 Science Center Dr, San Diego, CA 92121, CA. LSP Consulting LLC, Temecula, CA, 92591, USA. contributed equally to this work (Charlotte Bellamy, Hannah Tovell). contributed equally to this work (Marc Sanson, Alexandra C. Newton, Franck Bielle).

## Abstract

Chordoid glioma (ChG) is a rare, low-grade brain tumor characterized by a novel recurrent point mutation, D463H, in the kinase domain of protein kinase C alpha (PKCα). The mutation is invariably an Asp to His substitution, suggesting it endows a unique function beyond catalytic inactivation associated with other cancer-associated PKCα mutations. Here we use *in vitro* and *in cellulo* activity assays to show that PKCα_D463H_ is catalytically inactive, functions as a dominant-negative mutant to suppress endogenous PKC and uniquely rewires the cellular interactome. Specifically, phosphoproteomic, proximity labeling, and co-immunoprecipitation mass-spectrometry data from cells overexpressing PKCα_D463H_ identify altered phosphorylation of substrates and binding to multiple proteins involved in cell-cell junctions compared to WT enzyme. Lastly, single nuclei RNAseq reveals that ChG derives from specialized tanycytes. Our data suggest that this disease-defining, fully penetrant mutation promotes neomorphic non-catalytic scaffolding to impair cell junction function.

## Introduction

Protein kinase C alpha (PKCα) is mutated in diverse human cancers; however, highly penetrant disease-causing mutations are exceptionally rare^1^. A recurrent, fully-penetrant mutation has recently been identified in chordoid glioma (ChG), a rare, low-grade brain tumor ^2,3^. This mutation is the defining driver of ChG and involves the substitution of Asp463 in the highly conserved Y/HRD motif of the kinase domain^4^ to His. The invariant substitution to His specifically suggests a neomorphic function beyond simple loss-of-function and/or dominant-negative properties associated with the numerous other cancer-associated mutations that have been characterized in PKCα^5–7^. How this distinctive mutation alters PKCα is a unique paradigm for how neomorphic mutations might drive tissue-specific tumorigenesis. Identifying the mechanism of tumorigenesis of this unique PKC-driven cancer may inform on new approaches to PKC targeting therapies.

ChG is a rare, slow growing, brain tumor of grade 2 in the World Health Organization (WHO) classification^8^. It typically occurs in adults around the fifth decade, with a higher prevalence in females. ChG has a specific localization in the lamina terminalis at the anterior wall of the third ventricle, and is believed to be derived from tanycytes, specialized ependymal cells which line the third ventricle and contribute to the circumventricular organ (CVO). In this brain structure, the absence of blood-brain barrier enables the central nervous system to detect the blood levels of hormones and electrolytes. Tanycytes have the notable characteristic of being linked by tight junctions at their apical poles^9,10^, therefore establishing a functional barrier between permeable vessels and the third ventricle, unlike the rest of ependymal lining which lacks tight junctions in other brain structures where a blood-brain barrier is present^11^.

The ChG tumor itself is well circumscribed and nodular^12,13^. Histologically, it consists of clusters and cords of epithelioid cells, embedded in a mucin-like matrix with high infiltration by lymphocytes and plasmocytes^14^. Immunohistochemical studies indicate a glial lineage of the ventral forebrain^8,15,16^. This tumor context is particularly informative because the high recurrence of the D463H mutation and specificity for this rare tumor strongly suggests a unique cellular or tissular dependent mechanism. The main questions that arise are how the D463H mutation in PKCα functions at the cellular level, and why this mutation is restricted to this specific tissue and tumor.

PKCα is involved in multiple signaling pathways and regulates cellular functions such as growth, differentiation, cell adhesion, proliferation, and apoptosis^7^. PKCα signaling is finely tuned, and disturbances in PKCα substrate phosphorylation lead to pathologies. Whereas somatic loss-of-function mutations of PKC isozymes are generally associated with cancer^5,6^, rare and highly penetrant gain-of-function germline mutations are associated with Alzheimer’s Disease^17,18^. Loss-of-function mutations occur throughout the domain structure of PKC isozymes and either impair activation mechanisms to reduce activity or impair autoinhibitory constraints, which paradoxically results in loss-of-function as the active conformation of PKC is degradation-sensitive^19^. Despite the broad array of PKC mutations that have been associated with cancer, how rare driver mutations such as D463H then rewire PKC mechanisms in the cell and induce tissue-specific tumorigenesis, remains unknown.

PKCα belongs to the conventional subgroup of the nine-gene PKC family of Ser/Thr kinases. This subgroup is regulated by both diacylglycerol (DG), and Ca^2+ 20^. Newly translated PKCα protein is subject to a series of ordered phosphorylations by mTORC2 and PDK1 at specific motifs. These well described phosphorylations are necessary for the enzyme to adopt a catalytically-competent and autoinhibited conformation, through the binding of a pseudosubstrate that occupies the substrate-binding cavity^21–25^. This autoinhibited conformation is essential for the stability of the enzyme as the open, non-autoinhibited PKC is phosphatase labile, with the dephosphorylated species subject to ubiquitylation and degradation^26^. Binding of PKCα to its second messengers Ca^2+^ and DG results in membrane binding and release of the autoinhibitory pseudosubstrate to allow substrate binding and downstream signaling^27,28^.

Given this tightly regulated biology, understanding the biochemical and functional impact of the highly recurrent and unique to ChG mutation, PKCα D463H^2,3^, would provide the foundation to develop pharmacological treatments for the disease. Here we use biochemical, cellular, molecular dynamics, and proteomics approaches to gain insight into how the D463H mutation alters signaling and drives ChG tumorigenesis. Our results reveal a novel neomorphic non-kinase mechanism of PKCα. We show that the mutation renders PKCα catalytically inactive and that the overexpressed protein is strongly dominant-negative towards the activity of endogenous PKC isozymes. Molecular dynamics reveal that this mutation uniquely structures the substrate-binding regions of the C-lobe. Proximity biotinylation experiments reveal an enhanced interaction with 220 proteins. Our data are consistent with a model in which the D463H mutation uniquely binds many proteins involved in cell-cell junctions and cell polarity to rewire signaling events at these locations. This disruption in the context of the highly specialized tanycytes could initiate tumor formation by causing loss of tight-junctions and cell polarity.

## Results

### PKCα_D463H_ is an inactive kinase

PKCα D463 is a critical residue within the conserved Y/HRD motif of protein kinases, where it serves as the catalytic base to the phospho-acceptor residue of the substrate and helps with Mg^2+^ coordination^4^. Therefore, we first assessed whether PKCα_D463H_ can still catalyze substrate phosphorylation. GST-tagged PKCα_WT,_ PKCα_D463H_, or the previously characterized catalytically inactive PKCα_D463N_^29^ were isolated from Sf9 cells using a baculovirus expression system and the activity of purified proteins was assessed towards a peptide substrate [derived from substrate protein myristoylated alanine-rich C-kinase substrate (MARCKS)], either in the absence or presence of Ca^2+^, diacylglycerol (DG), and phosphatidylserine (PS). Purified GST-PKCα_WT_ robustly phosphorylated the peptide substrate in a Ca^2+^ and lipid-dependent manner, whereas neither GST-PKCα_D463H_ nor GST-PKCα_D463N_ had detectable kinase activity *in vitro* (Figure 1A).

**Figure 1.**
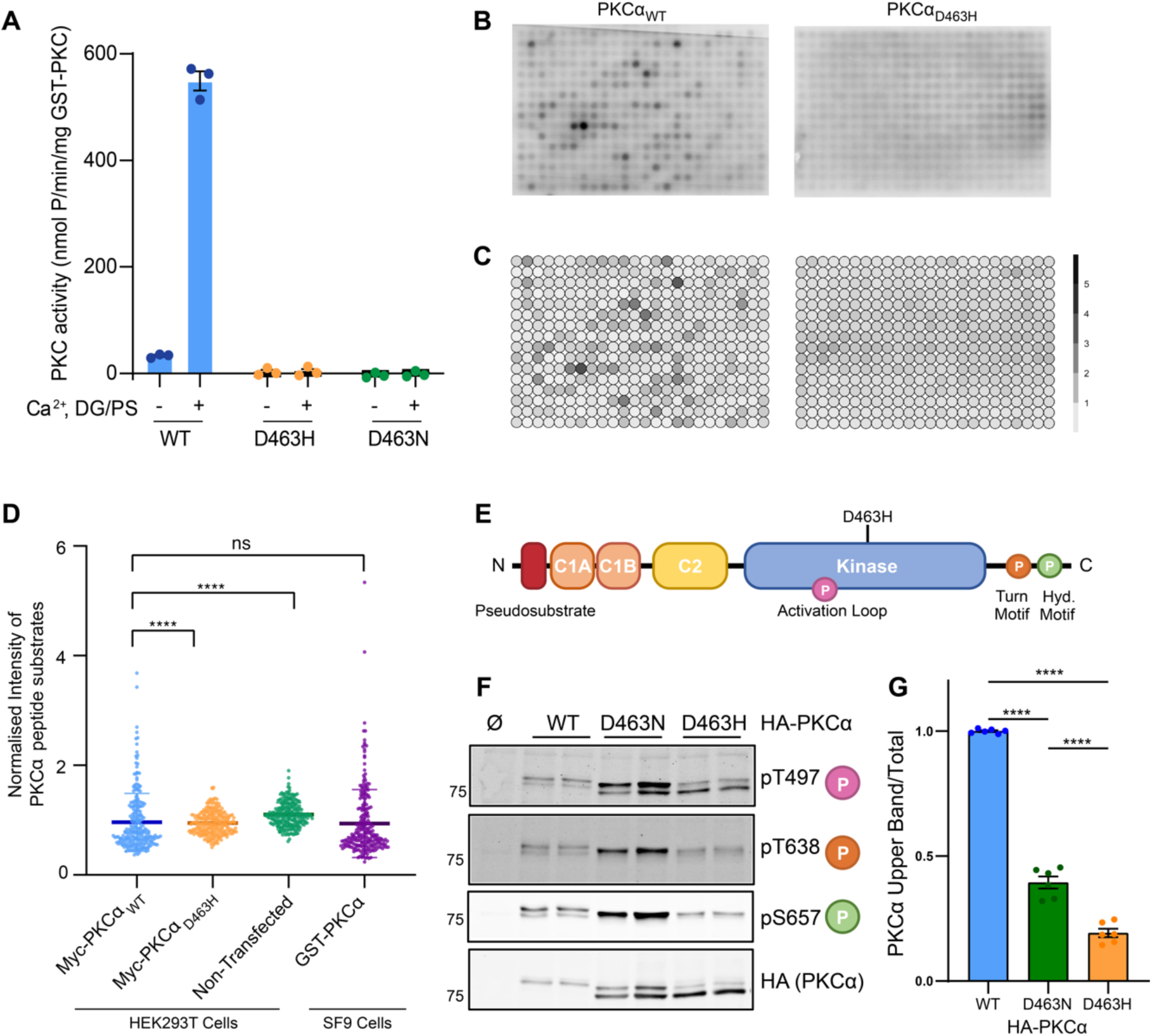
PKCα_D463H_ is an inactive kinase with a global loss of the phosphorylation of its known substrates. **A)** 20 ng GST-PKCα_WT_, GST-PKCα_D463H_, GST-PKCα_D463N_ was subject to *in vitro* kinase assay against MARCKS peptide substrate for 5 min in the presence or absence of activators DG/PS and Ca^2+^. **B)** Myc-PKCα_WT_ or Myc-PKCα_D463H_ were isolated by immunoprecipitation from HEK293T cells. Proteins were subject to kinase assay for 30 min across a peptide array of PKCα substrates, incubated with purified PKC, PKC activators DG/PS, Calcium and PMA, and ATP [γ-32P]. Array was imaged by autoradiography. **C)** Autoradiography intensities of peptide spots from D) normalized to scrambled peptide negative controls, n=3. **D)** Quantification of normalized autoradiography intensities from D), n=3, bar represents mean intensity across all replicates. (**** p < 0.0001, two-tailed Mann-Whitney test) **E)** Domain Structure of PKCα. PKCα is a member of the conventional PKC family, with 2 tandem DG-binding C1 domains (orange), and a Ca^2+^-binding C2 domain (yellow). Catalysis is performed by a kinase domain (blue) and regulated by an AGC C-tail and an N-terminal pseudosubstrate (red), which sits in the active site in the absence of agonist. Phosphorylation occurs at the activation loop (pink), turn motif (orange) and hydrophobic motif (green). **F)** COS7 cells were transfected for 48 h with HA-empty, or HA-PKCα constructs indicated, before lysis and probing for processing phosphosites by western blot. **G)** Quantification of HA-PKCα upper band intensity, relative to total HA-PKCα. Bars represent mean +/- SEM from n=3 experiments (**** p < 0.0001, 1-way ANOVA and Tukey multiple comparisons test).

To determine whether the D463H mutation in PKCα altered specificity towards substrates other than the MARCKS peptide, we interrogated kinase activity across a larger panel of substrates. Specifically, we designed a peptide array of 384 15mer peptides, representing known Ser/Thr substrates of PKCα and other kinases alongside negative controls (Table S1) and incubated this array with Myc-PKCα_WT_ and Myc-PKCα_D463H_, purified from HEK293T cells, ATP [γ-32P], PKC activators PS/DG, Phorbol 12-myristate 13-acetate (PMA), and Ca^2+^. Purified Myc-tagged PKCα_WT_ phosphorylated known substrates of PKCα with a similar profile to that of a commercially produced PKCα_WT_ (Figure S1A-B). In contrast, PKCα_D463H_ did not phosphorylate any of the diverse substrate peptides, confirming that the mutant protein is catalytically inactive (Figure 1B-D). Taken together, these experiments demonstrate that the amino acid substitution D463H in the active site of PKCα prevents kinase activity.

Retention of phosphate at the priming sites on PKCα serves as a marker of the autoinhibited conformation of the mature protein. Primed PKCα is quantitatively phosphorylated at two C-terminal sites, the turn motif (T638) and the hydrophobic motif (S657) (Figure 1E), with the latter phosphorylation promoting the autoinhibited and phosphatase-resistant conformation. To determine whether D463H impaired the autoinhibited conformation, we assessed the phosphorylation state of HA-tagged PKCα_WT_, PKCα_D463H_, and PKCα_D463N_ expressed in COS7 cells (Figure 1F). Phosphorylation at the C-terminal sites results in an electrophoretic mobility shift on SDS-PAGE gel^21^. Consistent with this, PKCα_WT_ migrated as a single slower mobility species, reflecting quantitative phosphorylation at the two C-terminal sites (Figure 1F, HA blot). As previously reported^29^, approximately 40% of D463N migrated as the slower mobilitiy species (Figure 1F and 1G) and this species was phosphorylated at all three priming sites (Figure 1F). In contrast, the majority (∼80%) of the D463H protein migrated as the faster-mobility species with the minor, slower mobility species (∼20%) phosphorylated at all three sites (Figure 1F and 1G). Despite mutation of the same residue, the reduced priming phosphorylations of the D463H compared to D463N revealed that the two mutants have differences in their conformation, with the D463H in a considerably more phosphatase-sensitive conformation.

### PKCα_D463H_ is dominant-negative and able to bind to PKCα_WT_

We next assessed the effect of expressing PKCα_D463H_ on the PKC signaling output of live cells. We measured the agonist-evoked activity of PKCα_WT_ and PKCα_D463H_ in cells using a C kinase activity reporter (CKAR2)^30^. Phosphorylation of this reporter results in a conformational change and increase in FRET signal^31^ that serves as a readout of PKC activity. COS7 cells expressing CKAR2 and mCherry (mCh) or mCh-PKCα WT or D463H were treated with 1) Uridine triphosphate (UTP), which transiently increases Ca^2+^ and DG to activate PKC^32^, 2) Phorbol 12,13-dibutyrate (PDBu), a non-metabolizable DG functional mimic which maximally activates PKCα^33,34^, and 3) Calyculin A (CalA)^35^, to suppress phosphatase activity and allow maximal phosphorylation of the reporter. The FRET/CFP ratio of the reporter was measured in real time through fluorescence microscopy (Figure 2A). In cells expressing CKAR2 and mCherry alone, UTP treatment resulted in a small, transient increase in PKC activity, which then returned to baseline within 5 min (Figure 2A, grey). This represents the activity of endogenous PKC isozymes. Treatment with PDBu resulted in a further sustained increase in CKAR2 signal, and inhibition of phosphatases by Calyculin A maximally phosphorylated the reporter. Co-expression of CKAR2 with mCh-PKCα_WT_ resulted in a much larger response to UTP, and rapid, phosphorylation of the reporter in response to PDBu (Figure 2A, blue). PDBu treatment maximally phosphorylated the reporter as no further increase was observed after Calyculin A treatment. This increase in CKAR2 signal relative to endogenous PKCs represents the activity of PKCα_WT_. Strikingly, expression of PKCα_D463H_ abolished the UTP-stimulated CKAR2 phosphorylation, revealing a strong dominant-negative effect of PKCα_D463H_ over endogenous PKC isozymes (Figure 2A, orange). In response to PDBu, CKAR2 phosphorylation increased at a rate and magnitude similar to mCherry alone, suggesting that saturating amounts of non-metabolizable agonist permit the activity of endogenous PKC isozymes. These results reveal that PKCα_D463H_ is strongly dominant-negative towards other PKC isozymes. This dominant-negative effect of PKCα_D463H_ was not observed against the related protein kinase A (PKA) (Figure S2A). The dominant-negative effect of PKCα_D463H_ towards other PKCs is particularly relevant as the PKCα D463H mutation is heterozygous in ChG, and all tumors retain one copy of *PRKCA* WT alongside a mutant copy^2^.

**Figure 2.**
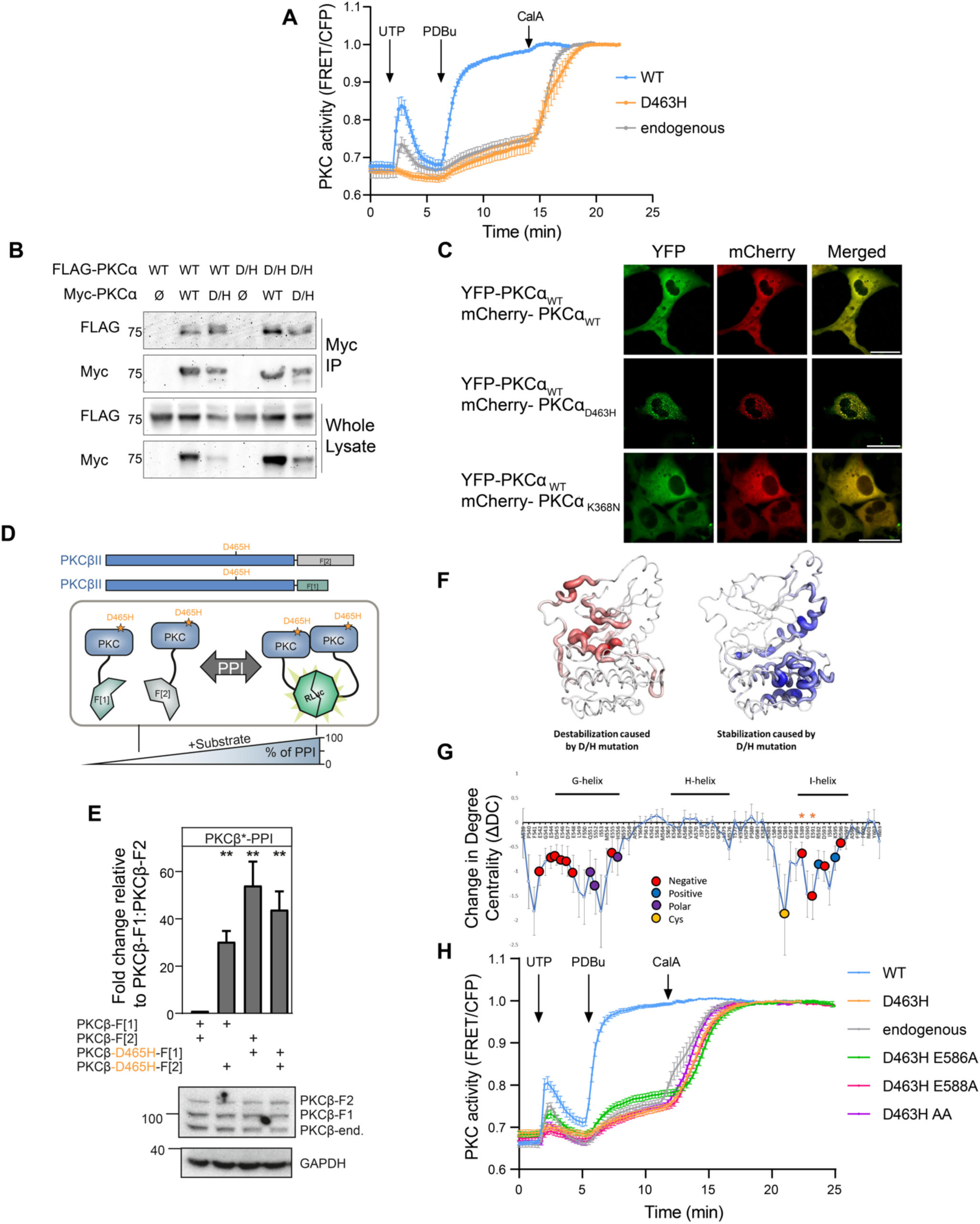
PKCα_D463H_ is dominant-negative and binds to PKCα_WT_ *in cellulo*. **A)** COS7 cells were transfected with CKAR2 PKC activity reporter and indicated mCherry-tagged PKCα constructs. 24 h post-transfection, cells were treated sequentially with UTP (100 µM), PDBu (200 nM), and Calyculin A (CalA; 50 nM). Data show FRET ratio changes normalized to maximum value, n=3 experiments and N≥30 individual cells, curve shows mean +/- SEM for each condition. **B)** HEK293T cells were co-transfected with Myc-PKCα WT or D463H, and FLAG-PKCα WT or D463H. 48 h post-transfection, cells were lysed and subject to immunoprecipitation with anti-c-Myc agarose beads. Eluted IPs were analysed by western blot for immunoprecipitated Myc-PKCα and co-immunoprecipitated FLAG-PKCα. **C)** COS7 cells grown on imaging slides were transfected with YFP-PKCα_WT_ and mCherry-PKCα WT, D463H, or K368N. 24 h post-transfection, cells were imaged by confocal microscopy. Scale bar: 30 µm. **D)** Mechanism of dimerization assay. PKCβII isoforms were tagged with PCA fragments F[1] and F[2] of the Renilla Luciferase. Dimerization *in cellulo* induces the complementation of fragments displaying bioluminescent signals reflecting quantifiable protein-protein interactions (PPIs). **E)** The indicated Rluc-based PCA reporter constructs were transiently co-expressed in HEK293T cells. Data are presented as the fold change in bioluminescent signals [relative light units (RLU)] relative to the signals of basal PKCβII homodimer formation. Bars indicate mean +/- SEM from five independent experiments. (** p <0.01, one sided Student’s *t* test) **F)** Molecular Dynamics modelling of the stability of PKCβII WT and D465H mutant, with stability assessed using LSP alignment. Left – regions where the WT shows greater stability compared to the D465H mutant (positive ΔDC values, shown in red). Right – regions where the mutant shows greater stability compared to the WT (negative ΔDC values, shown in blue). **G)** Change in Degree Centrality (ΔDC) demonstrating increased stability of the D465H mutant protein, focusing on the G-H-I helices of the kinase C-lobe. Error bars represent the standard error (SE) calculated from five replicate simulations. The residues are color coded by type. The two residues marked with orange asterisks were selected for further mutation in panel H. **H)** COS7 cells were transfected with CKAR2 PKC activity reporter and indicated mCherry-tagged PKCα constructs. 24 h post-transfection, cells were treated sequentially with UTP (100 µM), PDBu (200 nM), and Calyculin A (50 nM). Data show FRET ratio changes normalized to maximum value, n=3 experiments and N≥30 individual cells, curve shows mean +/- SEM for each condition.

We next used a co-immunoprecipitation approach to understand whether the dominant-negative effect of this mutation results from a physical interaction of PKCα_D463H_ with PKCα_WT_. FLAG and Myc-tagged PKCα WT or D463H mutant, were co-expressed in HEK293T cells and isolated by Myc-immunoprecipitation, followed by western blot for co-immunoprecipitated FLAG-tagged PKCα. Myc-PKCα_WT_ co-immunoprecipitated a faster mobility (unphosphorylated) species of FLAG-PKCα_WT_ as previously described^25^ and also immunoprecipitated FLAG-PKCα_D463H_ with substantially greater efficiency (Figure 2B, compare lanes 2 and 5). Consistent with enhanced binding resulting from the mutation, Myc-PKCα_D463H_ was more effective than WT at immunoprecipitating FLAG-PKCα_WT_ (Figure 2B, compare lanes 2 and 3). This interaction between PKCα_WT_ and PKCα_D463H_ could be a potential mechanism by which the mutant protein is dominant-negative over WT.

The D463H mutation abolishes catalytic activity of PKCα, yet other *PRKCA* mutations that also inactivate PKCα are not observed in ChG. This suggests that the amino acid change is not simply a loss-of-function mutation. To explore the functional effects specific to His at position 463, we asked whether D463H, but not other kinase dead constructs, altered the subcellular distribution of PKCα_WT_. We therefore monitored the subcellular localization of YFP-PKCα_WT_ co-expressed with either mCh-PKCα_WT_, PKCα_D463H,_ or the kinase-inactive PKCα_K368N_ in COS7 cells (Figure 2C). YFP-PKCα_WT_ displayed diffuse cytosolic staining in the presence of mCh-PKCα_WT_. Strikingly, co-expression with PKCα_D463H_, but not PKCα_K368N_, resulted in redistribution of the WT enzyme to puncta throughout the cell (Figure 2C). PKCα_D463H_ alone also displayed this distribution to puncta (Figure S2B). The sequestration of PKCα_WT_ away from its endogenous targets to potentially new interactors provides an attractive mechanism for the unique effects of His at position 463 beyond those observed with other inactivating mutations.

We previously employed a luciferase-based protein-fragment complementation assay (PCA) to quantify PKCβII protein dimerization in intact cells^25^ (Figure 2D). As PKCβII and PKCα are highly paralogous (Figure S2C), with 80% sequence homology and high structural similarity, we used this system to examine the influence of the ChG mutation (D465H in PKCβII) on PKCβII dimerization within a cellular context. Utilizing this approach, we observed a significant enhancement of PKCβII dimerization upon integration of the D465H mutation into either protomer (Figure 2E). Intriguingly, the D465H mutation also increased the very weak affinities of PKCβII for PKA RIα (Figure S2D). Building on evidence that mutation of Asp to His in the Y/HRD motif imparts unique properties to PKC, we investigated how this mutation affects thermal motions within the kinase domain through computational methods. Using molecular dynamics (MD) simulations and Local Spatial Pattern (LSP) alignment, we analyzed the impact of the D465H mutation on sub-nanosecond dynamics in the PKCβII kinase domain. These fast dynamics play an essential role in protein function and molecular recognition, influencing protein-ligand and protein-protein interactions by modulating conformational entropy^36^. LSP alignment is a graph-theory-based approach that quantifies CαCβ vector mobility in molecular dynamics trajectories. From a 10 nanosecond simulation, we created a graph where residues are nodes and edge weights represent their relative stability. This approach has been shown to capture changes in conformational entropy and reveal long-distance allosteric effects following ligand binding^37^. Here, we applied LSP alignment to study the allosteric effects of the ChG mutation in the PKCβII kinase domain. Following previous methods, we summed the weights of each residue’s edges to calculate Degree Centrality (DC). Since the weights estimate stability, DC serves as an indicator of local stability, with high DC values reflecting more stable regions and low DC values suggesting higher mobility.

Modelling was performed using the well-characterized solved structure for the kinase domain of PKCβII (PDB: 2I0E)^25,38,39^. Given the high sequence similarity between PKCα and PKCβII, the PKCβII structure serves as a good approximation for our analysis. LSP analysis identified several regions with predicted altered stability between the WT and D465H PKCβII kinase domain. The ATP-binding pocket of the kinase domain is predicted to be significantly disrupted by the D465H mutation, with greater dynamic stability observed in the WT model (Figure 2F, left panel). This outcome is expected as D465 not only acts as a proton acceptor for the substrate but also plays a critical role in coordinating the Mg^2+^ ion within the active site, allowing ATP binding. Surprisingly, our model also predicted significant changes in dynamics in regions distant from the mutated residue. Notably, we observed an increase in stability within the G-, H-, and I-helices of the C-lobe in the mutant model (Figure 2F, right panel). Figure 2G demonstrates the change in Degree Centrality (ΔDC) of this region, where ΔDC represents the difference in stability between the WT and the mutant models: positive ΔDC values indicate greater stability in the WT, while negative values reflect increased stability in the mutant. Here, the G- and I-helices show higher DC in the mutant, and therefore increased stability. The I-helix of the kinase domain has previously been associated with binding of protein substrates^40,41^. These results support a model in which altered dynamics in the substrate-binding surface of the C-lobe promotes not only increased binding to PKC substrates but also potentially aberrant binding to unnatural partners.

To test the hypothesis that alterations in the I-helix drive the unique properties of His instead of Asp in the Y/HRD motif, we asked whether mutating the surface of the I-helix to reduce substrate binding would reduce the dominant-negative effect of PKCα_D463H_ as assessed using CKAR2. We combined the D463H (D465H in PKCβII) mutation with two mutations in the I-helix of the kinase – E586A (E589 in PKCβII) and E588A (E591 in PKCβII), demonstrated with an orange asterisk in Figure 2G. These mutations were selected because E591 was the most stabilized residue within the I-helix of the kinase domain, and disruption of the negative charge on these residues was predicted to impact protein binding. As observed in Figure 2A, expression of mCh-PKCα_D463H_ suppressed the UTP-stimulated activity of endogenous PKC in COS7 cells (Figure 2H, orange). Expression of PKCα_D463H/E586A_ effectively rescued this dominant-negative effect, restoring endogenous UTP-stimulated activity (Figure 2H, green). The PKCα_D463H/E588A_ and PKCα_D463H/E586A/E588A_ constructs did not restore activity (Figure 2H, pink and purple traces). These data reveal that alterations in the I-helix contribute to the dominant-negative phenotype of PKCα_D463H_.

### PKCα_D463H_ has a more open and less stable conformation

PKCα_D463H_ has previously been shown to be more proteolytically unstable than the WT enzyme^2^. As PKC_WT_ is typically held in a stable form by occupancy of the active site by the pseudosubstrate, we explored whether this instability was related to decreased occupancy of the mutant active site. We therefore compared the effect of the D463H mutation to the pseudosubstrate mutation A25E, previously shown to promote the proteolytically labile conformation of PKC and predicted to be proximal to D463 in the PKCα active site^39,42^. Additionally, the entire pseudosubstrate (PS), amino acids 2-29^43^, was deleted (ΔPS) in both PKCα_WT_ and PKCα_D463H_ (Figure 3A). COS7 cells expressing these constructs were treated with cycloheximide (CHX), which inhibits protein synthesis and allows for the assessment of protein stability, for up to 48 hours, and the rate of PKCα degradation measured by western blot (Figure 3B, 3C). In agreement with previous data, the properly autoinhibited PKCα_WT_ had a half-life greater than 48 hours (Figure 3B, blue). In contrast, the half-life of PKCα_D463H_ dramatically decreased (t_1/2_ = 16 ± 5 h). The unphosphorylated (faster mobility), predominant species was rapidly degraded, whereas the slower mobility phosphorylated species was degraded at a comparable rate to the wild-type enzyme in the first 8 hours (Figure S3A). The degradation rate of PKCα_D463H_ was similar to that of PKCα_A25E_ and PKCα_ΔPS_ (Figure 3B, 3C). Importantly, combination of pseudosubstrate deletion with the D463H mutation caused a nearly four-fold decrease in half-life (PKCα_ΔPS/D463H_ t_1/2_ = 4.1 ± 0.8 h) compared to D463H alone, indicating that disengagement of the pseudosubstrate from the active site is not the sole mediator of PKCα_D463H_ instability (Figure 3B, 3C).

**Figure 3.**
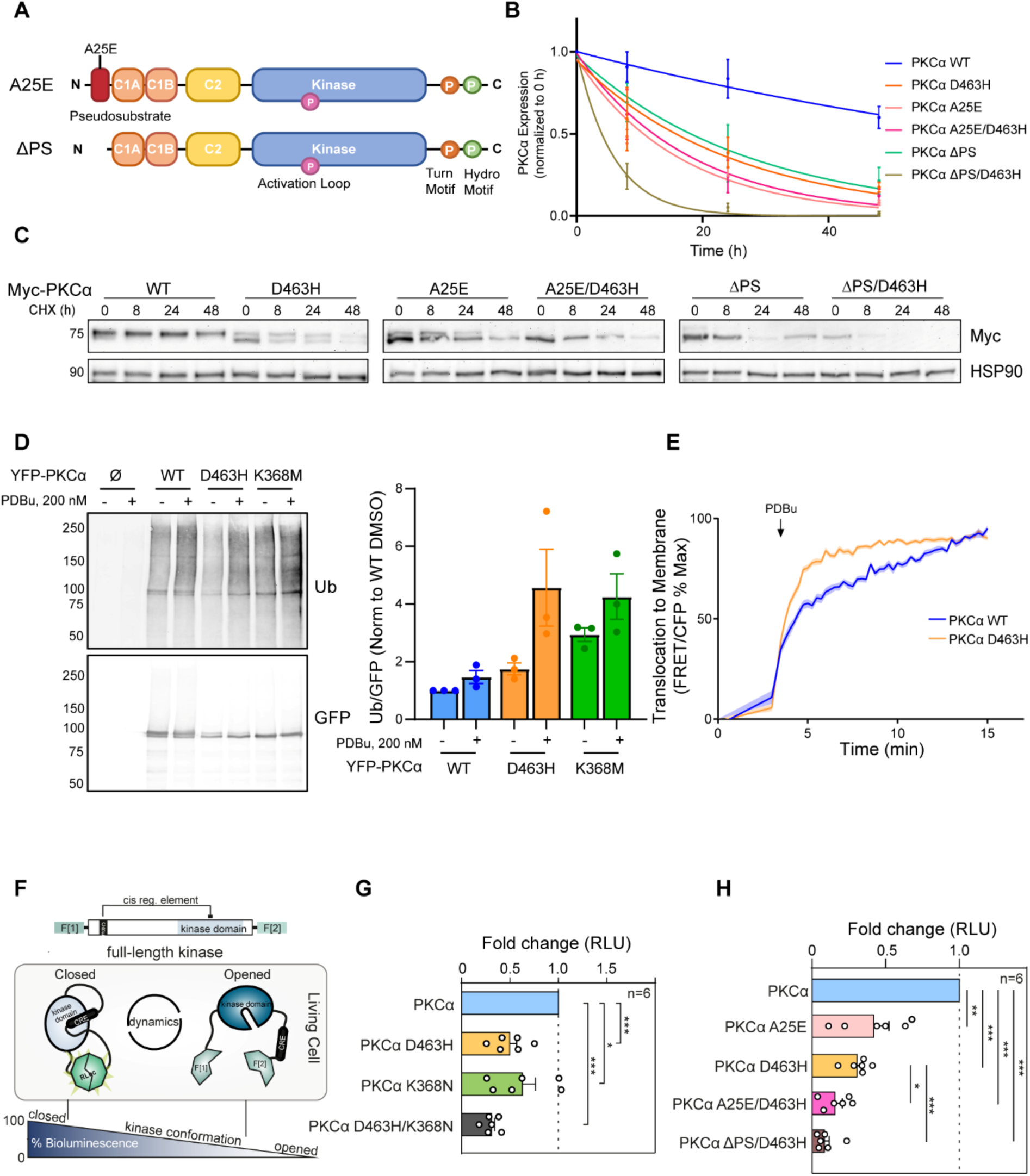
PKCα_D463H_ has a more opened and less stable conformation. **A)** Domain structure of PKCα_A25E_ constructs, carrying pseudosubstrate A25E mutation, and PKCα_ΔPS_, carrying deletion of residues 2-29. **B)** Quantification of Myc-PKCα protein levels from C), Myc western blot intensity normalized to loading control and 0 h timepoint. Data were fit to first order decay, curve represents mean +/- SEM, n=3. **C)** Indicated PKCα constructs were overexpressed in COS7 cells for 24 h before treatment with 355 µM cycloheximide (CHX) for the times indicated. Cells were lysed and subject to western blot to assess Myc-PKCα protein levels. **D)** YFP-empty (Ø) or indicated YFP-PKCα constructs were overexpressed in COS7 cells. 48 h post-transfection, cells were treated for 8 h with 20 µM MG132 to prevent proteasomal degradation and either DMSO or 200 nM PDBu for the final 4 h. Cells were lysed, YFP-PKCα was isolated by immunoprecipitation and ubiquitination levels measured by western blot. **E)** Indicated YFP-PKCα constructs and MyrPalm-CFP were overexpressed in COS7 cells for 24 h. Cells were treated with 200 nM PDBu and translocation of PKCα to plasma membrane was measured by quantification of FRET by live fluorescent microscopy. Graph shows FRET/CFP ratios, normalized to maximum FRET/CFP for each condition, n=4 and N≥30 cells per condition, curves show mean +/- SEM for each condition. **F)** Schematic depiction of the Kinase Conformation (KinCon) reporter principle for measuring auto-inhibitory kinase conformations (cis regulatory elements) in living cells. Indicated PKCα constructs were flanked with PCA fragments of the Renilla Luciferase (RLuc). Autoinhibited PKCα are indicated through RLuc protein-fragment complementation and higher bioluminescence signals whereas a PKCα with reduced autoinhibition generates reduced luciferase activity. **G)** Fold change of the KinCon reporter signals from the PKCα KinCon reporters are displayed for the indicated mutations relative to the WT PKCα reporter values (n=6 individual experiments, normalized to KinCon reporter expression levels, mean +/- SEM; significance was determined by Welch’s t test with * p <0.05, ** p <0.01, *** p <0.001). **H)** As in G, analysis of the PKCα KinCon reporters are displayed for the pseudosubstrate mutations indicated.

To assess whether the faster rate of PKCα_D463H_ degradation resulted from increased ubiquitination of the protein, we compared the basal and agonist-induced ubiquitination of PKCα_WT_, PKCα_D463H_, and kinase-inactive PKCα_K368M_. COS7 cells expressing each of these constructs were treated with proteosome inhibitor MG132 (20 µM) to accumulate ubiquitinated protein in the presence or absence of PDBu. Analysis of ubiquitination levels by YFP-immunoprecipitation and western blot revealed that all PKCα proteins were basally ubiquitinated and accumulated additional ubiquitination after treatment with PDBu. The basal ubiquitination of PKCα_D463H_ was modestly increased (1.5-fold higher) in comparison to PKCα_WT_, whereas PKCα_K368M_ ubiquitination was substantially higher (∼3-fold higher) (Figure 3D). PDBu treatment resulted in a modest increase (<2-fold) in ubiquitination of the WT, whereas PKCα_D463H_ and PKCα_K368M_ both displayed 3-fold increases in ubiquitination compared to WT. The differing basal ubiquitination of D463H compared to K368M suggests these mutants adopt different conformations within the cell, although both are rapidly ubiquitinated in response to PDBu (Figure 3D).

We next used FRET to quantify the kinetics of PDBu-induced translocation of YFP-tagged PKCα_WT_ or PKCα_D463H_ towards plasma membrane-targeted CFP^19,39,44^. We have previously shown that a more open PKCα that is not in an autoinhibited conformation translocates to the membrane at a faster rate^19^. As observed previously, PKCα_WT_ translocated to the membrane with biphasic kinetics, whereas PKCα_D463H_ abolished the slower phase, a phenomenon previously reported for K368M and D463N^19^, therefore translocating at a much faster rate (Figure 3E, Figure S3B). These data are consistent with PKCα_D463H_ adopting an open conformation.

To confirm that the reduced stability, increased ubiquitination, and increased membrane translocation resulted from a less autoinhibited conformation of PKCα_D463H_ compared to PKCα_WT_, we utilized an intramolecular based protein-fragment complementation assay^45,46^ (termed KinCon reporter) to assess PKCα conformations in intact cells^25,47^. PKCα_WT_ and PKCα_D463H_ constructs flanked with the N-terminal and C-terminal fragments of luciferase were expressed in COS7 cells and luciferase activity was used as a measure of the degree of ‘closed’ conformation (Figure 3F). Relative to PKCα_WT_, PKCα_D463H_ KinCon reporter protein was in a more opened conformation, as assessed by the ∼50% reduction in luciferase signal (Figure 3G). We additionally combined these reporters with the K368N kinase dead mutation, and the pseudosubstrate mutants. The PKCα_K368N_ based KinCon reporter, which is known to lack autoinhibition, produced a similar reduction in signal compared to PKCα_D463H_ (Figure 3G). Notably, combination of these two mutations (PKCα_D463H/K368N_) resulted in a further reduction in luciferase signal (∼75% reduction in comparison to PKCα_WT_), suggesting that both mutations synergize to further promote the opening of cellular PKCα conformations and reduce autoinhibition (Figure 3G). Integration of the pseudosubstrate mutation resulted in opening of the PKCα_A25E_ KinCon reporter, as expected, and PKCα_A25E/D463H_ combined to further open the reporter. (Figure 3H). This is in concordance with the cycloheximide data where a more open conformation results in more rapid degradation. Deletion of the pseudosubstrate combined with the D463H mutation (PKCα_ΔPS/D463H_) caused the largest decrease in luciferase signal, reflecting the most efficient opening of the KinCon reporter conformation of PKC (Figure 3H). The expression levels of the KinCon reporter constructs are shown in Figure S3C.

### PKCα_D463H_ dysregulates signaling pathways involved in cell junctions

PKCα regulates a vast array of pathways, impacting on many cellular functions such as cell proliferation, survival and cytoskeletal maintenance^7^. To understand the effect the mutation had on global protein phosphorylation, we undertook phosphoproteomic mass-spectrometry analysis of HEK293T cells overexpressing either FLAG-PKCα_WT_ or FLAG-PKCα_D463H_. This analysis revealed a dramatic decrease in protein phosphorylation in cells overexpressing the mutant. We observed 40 phosphosites decreased in the mutant condition and 66 phosphosites uniquely present upon expression of PKCα_WT_ (fold change (FC)>1.5, p<0.05). Upon expression of PKCα_D463H_, we observed only 3 phosphosites increased and 3 phosphosites uniquely present (FC>1.5, p<0.05) (Figure 4A and Table S2). These changes were not due to alterations at the protein level as determined by parallel proteome analysis (Figure S4A). Kinase activity profiles determined from the phosphoproteomic data using Integrative iNferred Kinase Activity (INKA) analysis of altered phosphosites^48^ confirmed a reduction in PKCα activity (Figure S4B). Gene ontology (GO) enrichment analysis^49,50^ for proteins with significantly decreased phosphorylation sites (FC>1.5, p<0.05) identified several pathways impacted by PKCα_D463H_ expression. These include Cell Adhesion Molecule Binding (GO: 0050839, adjusted p-value=5.78e-03) and Cadherin Binding (GO: 0045296, adjusted p-value=1.69e-04) (Figure 4B). Amongst the proteins that had significantly reduced phosphorylation, we identified cell adhesion regulator protein Catenin Delta-1 (CTNND1), whose phosphorylation at Ser879 was reduced by 2.24-fold (adjusted p-value=0.02) (Figure 4A). This site has been previously described as a target of PKCα, and phosphorylation of this site is a known phosphoswitch leading to VE-Cadherin junction disassembly^51,52^. Another identified known phosphoswitch with a significant reduction in phosphorylation was Ser284 of Junctional Adhesion Molecule-1 (JAM-A) (3.07-fold, adjusted p-value=5.73e-03), a protein implicated in tight junction formation (Figure 4A). Phosphorylation of this site has been shown to be important for the maintenance and integrity of tight junctions^53^, which is of particular interest considering the importance of tight junctions in tanycyte function^10,54^.

**Figure 4.**
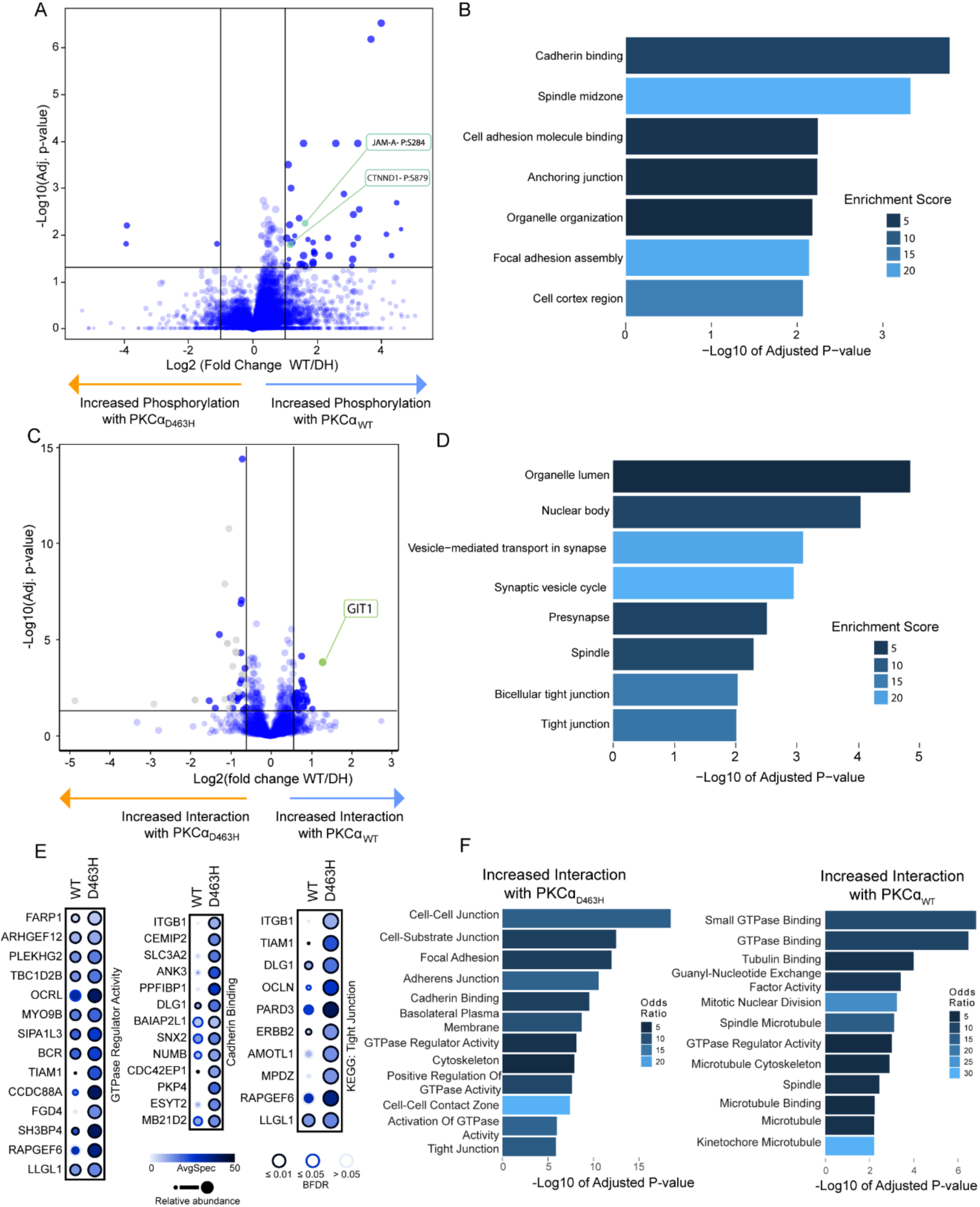
PKCα_D463H_ dysregulates signaling pathways involved in cell junctions. **A)** Phosphopeptide analysis of HEK293T cells overexpressing FLAG-PKCα_WT_ or PKCα_D463H_. HEK293T cells were transfected 48 h prior to lysis, followed by Trypsin/LysC digest and TiO_2_ phosphopeptide enrichment. n=5 samples per condition. CTNND1 and JAM-A phosphoswitches are highlighted. Significance threshold fold change (FC)>1.5, p<0.05. **B)** GO term enrichment analysis of proteins with a decrease in protein phosphorylation upon overexpression of PKCα_D463H_. **C)** Co-immunoprecipitation mass-spectrometry analysis of FLAG-PKCα_WT_ and PKCα_D463H._ HEK293T cells were transfected 48 h prior to lysis, followed by FLAG immunoprecipitation. Coimmunoprecipitating peptides were subject to Trypsin/LysC digest and quantified by mass-spectrometry. Significance threshold FC>1.5, p<0.05. **D)** GO term enrichment analysis of proteins that have significantly decreased interaction with PKC⍺_D463H_. **E)** Proximity Biotinylation analysis of HEK293T cells overexpressing FLAG-MiniTurbo-PKCα_WT_ and PKCα_D463H._ MiniTurbo-PKCα constructs were expressed in Flp-In™ T-REx™ 293 cells for 24 h before stimulation with biotin, cell lysis, and isolation of proximal proteins by Streptavidin pulldown and mass-spectrometry. Representative dotplots show proteins with preferred proximity to PKCα_D463H_ protein. Dot size represents change in spectral intensity, color represents number of spectra identified, p<0.05 represented by black outline. n=2 per condition. **F)** GO term enrichment analysis for proteins found to preferentially interact with PKCα_WT_ or PKCα_D463H_ by MiniTurboID analysis.

To explore further how the D463H mutation alters cellular function of PKCα, we performed co-immunoprecipitation mass-spectrometry (coIP-MS) to identify differential interactors of PKCα_WT_ and PKCα_D463H_ (Figure 4C). FLAG-PKCα_WT_ or FLAG-PKCα_D463H_ were overexpressed in HEK293T cells for 24 hours before performing FLAG immunoprecipitation, with co-immunoprecipitating interactors analyzed by mass-spectrometry. We detected 98 proteins that significantly lost interaction and 107 proteins that significantly gained interaction with the mutant protein (FC>1.5, p<0.05) (Figure 4C and Table S3). GO analysis of the proteins that displayed a loss of interaction with PKCα_D463H_, indicated enrichment for Tight Junctions (GO: GO:0070160, adjusted p-value=9.71e-0.3) (Figure 4D).

To better understand the interactome of PKCα_D463H_, we performed a biotin-mediated proximity labeling assay to determine dynamic interacting proteins in live cells. Flp-In™ T-REx™ 293 cells expressing Flag-MiniTurbo-PKCα_WT_ or Flag-MiniTurbo-PKCα_D463H_ were treated with doxycycline for 24 hours to induce PKCα expression before inducing biotinylation of interacting proteins by addition of exogenous biotin. It is important to note that the protein affinities that are being measured in the MiniTurbo assay and coIP-MS analysis are different levels of interactions. CoIP-MS analysis identifies very strong, stable interactions often observed in protein complexes while the MiniTurbo assay identifies temporal changes in proximity^55^.

We identified 220 proteins with preferred proximity to PKCα_D463H_ and 135 with enhanced proximity to PKCα_WT_. Proteins with preferential proximity to PKCα_D463H_ were enriched for GO Terms included: Cell-Cell Junctions (GO:0005911, adjusted p-value=3.38e-19, Cadherin Binding (GO:0045296, adjusted p-value=2.89e-10), and GTPase Regulator Activity (GO:0030695, adjusted p-value=7.60e-09 (Figure 4E and 4F, left). We identified several regulators of the Rho GTPases Rac1 and CDC42, which are known to regulate tight junctions and apical-basal polarity (Figure 4E, 4F and Figure S4C). These are pathways that are known to be regulated by PKCα and are particularly interesting in the context of the highly specialized, polarized tanycytes. Of the proteins with reduced interaction with the mutant and preferential proximity to the WT protein, enriched GO Terms included Mitotic Nuclear Division (GO:0140014, adjusted p-value=5.79e-04), and Spindle Microtubule (GO:0005876, adjusted p-value=7.81e-04), alongside Small GTPase Binding (GO:0031267, adjusted p-value=9.56e-10) (Figure 4F, right).

### PKCα_D463H_ alters cell-cell junction pathways in ChG

Next, we sought to better characterize molecular changes in the ChG tumor cells. ChG tumors contain a high immune infiltration, and therefore, to investigate pathways that are specifically altered in the tumor cells, we performed a single nuclei (snRNAseq) analysis of a ChG tumor sample and obtained 3112 nuclei after quality control (Figure 5A). We identified a high content of immune cells (n=999, 32%) made up of four distinct populations (plasmocytes, macrophages, lymphocytes, and microglia), as previously observed in tumor tissue samples^16^. Immunohistochemical studies have previously shown that ChG tumors express Glial fibrillary acidic protein (GFAP), vimentin, Thyroid transcription factor 1 (NKX2-1/TTF1), and CD34^13,15,16^. In the snRNAseq analysis, we identified four populations of tumor cells that shared expression of the ChG marker *NKX2-1/TTF-1*, but had distinct expression levels of *CD34, GFAP,* and the tyrosine kinase *NTRK3.* Using differentially expressed genes specific to each cluster, we performed a deconvolution analysis of bulk RNAseq data from nine different patient samples (Figure 5B). This analysis revealed that ChG tumors are predominantly made up of the cluster ‘*NTRK3* low, *CD34* high’ tumor cells (65-90% of cells) alongside a rich infiltration by immune cells (35-60% of cells) (Figure 5B). This analysis suggests that, despite the low number of samples available due to the rareness of this tumor, the snRNAseq data are most likely representative of all ChG tumors.

**Figure 5.**
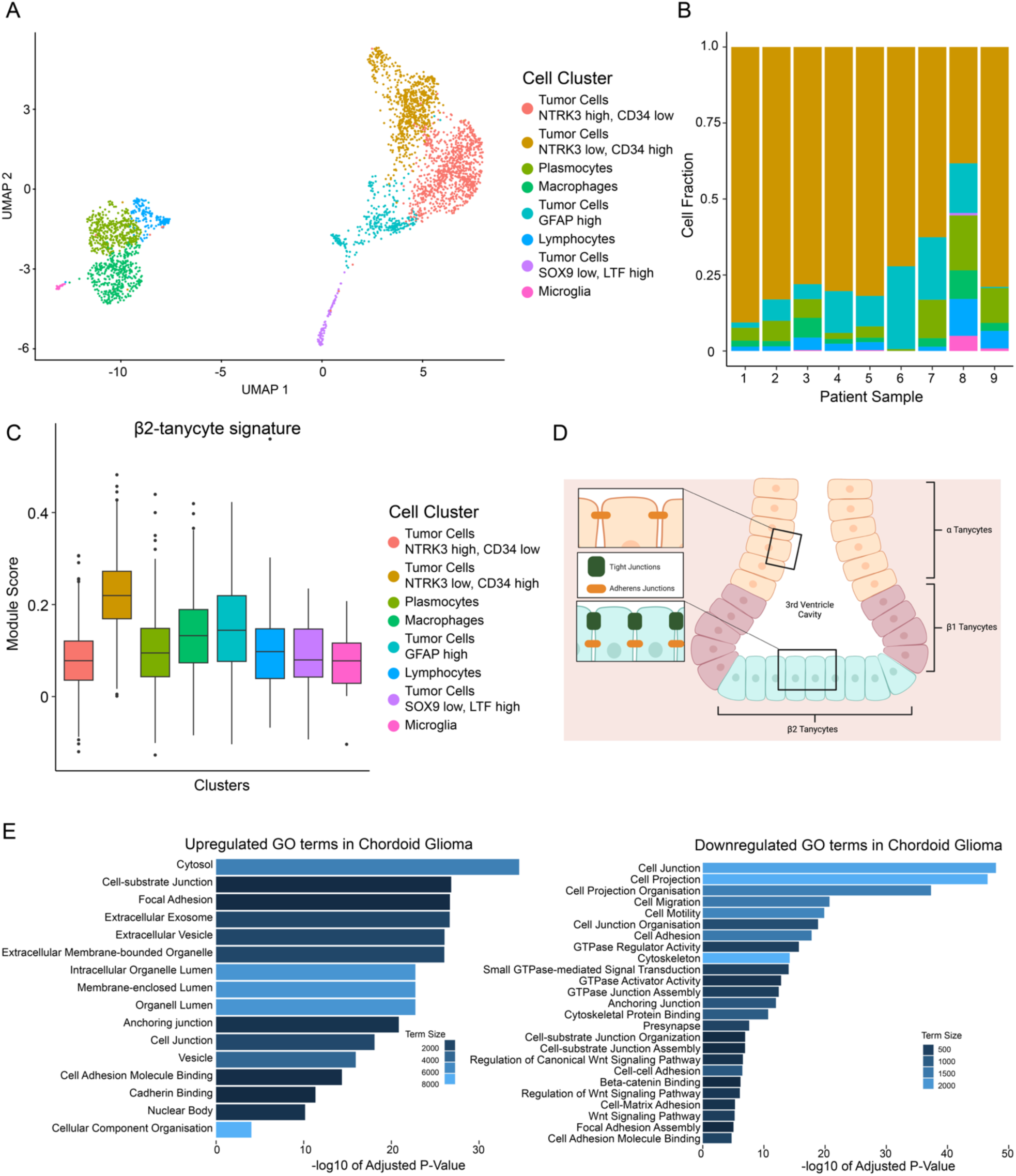
Chordoid gliomas develop from β2-tanycytes, with highly specialized tight junctions. **A)** UMAP plot of snRNAseq of a chordoid glioma sample, showing the different cellular clusters representing distinct cell populations. **B)** Deconvolution analysis of Bulk RNAseq data of ChG samples from 9 patients using snRNAseq data in A, showing the cell populations making up each sample. Coloring of cell populations as in A. **C)** Module score of β2-tanycyte signature (upregulated genes compared to other tanycyte subtypes) in the different cellular clusters of the ChG snRNAseq from A. **D)** Diagram showing location of tanycyte subtypes within the 3^rd^ Ventricle. Inset - junctions found between β2 or α tanycytes. **E)** Graphs show the most significant ontologies identified from GO enrichment analysis of the differentially expressed RNA that is (Left) upregulated in the ChG tumor cluster (*NTRK* low/*CD34* high) and (Right) downregulated in the ChG tumor cluster (*NTRK* low/*CD34* high) compared to healthy human astrocytes.

Histological studies have already suggested that ChG derives from a type of ependymal cell called tanycytes^16^, so we compared gene signatures from various sub-populations of tanycytes^56^ with the tumoral data. We found an enrichment for the β2-tanycyte signature within the predominant ‘*NTRK3* low and *CD34* high’ tumor population of ChG (Figure 5C). This cluster did not show enrichment for α- or β1-tanycyte signatures. Interestingly, β2-tanycytes specifically are connected by tight junctions (in comparison with other tanycytes) to produce a barrier at this specific location of the brain^57^ (Figure 5D). Differential analysis of the predominant tumor cluster ‘*NTRK3* low, *CD34* High’ with human normal astrocytes revealed differentially expressed genes in ChG that are enriched forseveral GO terms(Figure 5E). These altered gene sets are consistent with the GO terms identified through the proteomic analyses observed in HEK293T cells, including Cell Adhesion Molecule Binding (GO:0050839, adjusted p-value= 4.38e-15), Cadherin Binding (GO:0045296, adjusted p-value=4.61e-12), Focal Adhesion Assembly (GO:0048041, adjusted p-value=9.68e-06), and Anchoring Junctions (GO:0070161, adjusted p-value= 1.10e-12) (Figure 5E). Taken together with the global phosphoproteome and interactome analysis, dysregulation of pathways regulating cell junctions, including tight junctions, are the primary consequence of PKCα_D463H_ mutation. This is of particular interest given the importance of tight junctions in the cell of origin.

### PKCα_D463H_ alters cell junction protein localization in HEK293T cells and chordoid glioma

Our phosphoproteomic analysis identified phosphoswitches modified by the D463H mutant on proteins that regulate tight junctions: CTNND1 pSer879 and JAM-A pSer284. To understand the functional effect of these altered phosphoswitches, we overexpressed either YFP-PKCα_D463H_ or YFP-PKCα_WT_ in HEK293T cells and quantified the junctional markers by immunofluorescence using an analysis pipeline (Figure S5A-B). The phosphorylation at S879 on CTNND1 has been previously described as a destabilizing phosphorylation^52^. Phosphoproteomics identified a loss of phosphorylation at this site upon overexpression of PKCα_D463H_. We hypothesized that this loss would lead to stabilization of CTNND1 and therefore a higher level at cell-cell contacts. Upon quantification of this protein intensity at cell membranes between two contiguous YFP positive cells (meaning both were expressing exogenous PKCα), we saw a significant increase of intensity of CTNND1 in HEK293T cells expressing YFP-PKCα_D463H_ in comparison to YFP-PKCα_WT_ (Figure 6A, 6C). Immunohistochemistry (IHC) of ChG tumoral tissue also showed positivity for CTNND1 at cell junctions (Figure 6I, top) at a similar level to ependymocytes^11^, another type of brain epithelial glial cell known to connect by adherens junctions (Figure 6I, bottom). Due to our hypothesis of CTNND1 stabilization from loss of phosphorylation, we expected a concomitant increase of cadherin at these cell-cell contacts. However, we observed a significant decrease in the amount of N-Cadherin/CDH2, the cadherin found in neural cells^58^, at cell-cell contacts upon expression of YFP-PKCα_D463H_ (Figure 6B, 6D). This could be due to another cadherin mediating these junctions in ChG. Phosphorylation of JAM-A on Ser284 was also reduced in the phosphoproteomic analysis. Loss of this phosphorylation site has been reported to lead to dissociation of JAM-A from the tight junction complex, affecting the barrier function while not regulating the localization of JAM-A^53^. At cell-cell contacts between two transfected cells, a slight increase in JAM-A intensity was detected upon expression of YFP-PKCα_D463H_ compared to YFP-PKCα_WT_ (Figure 6E, 6G).

**Figure 6.**
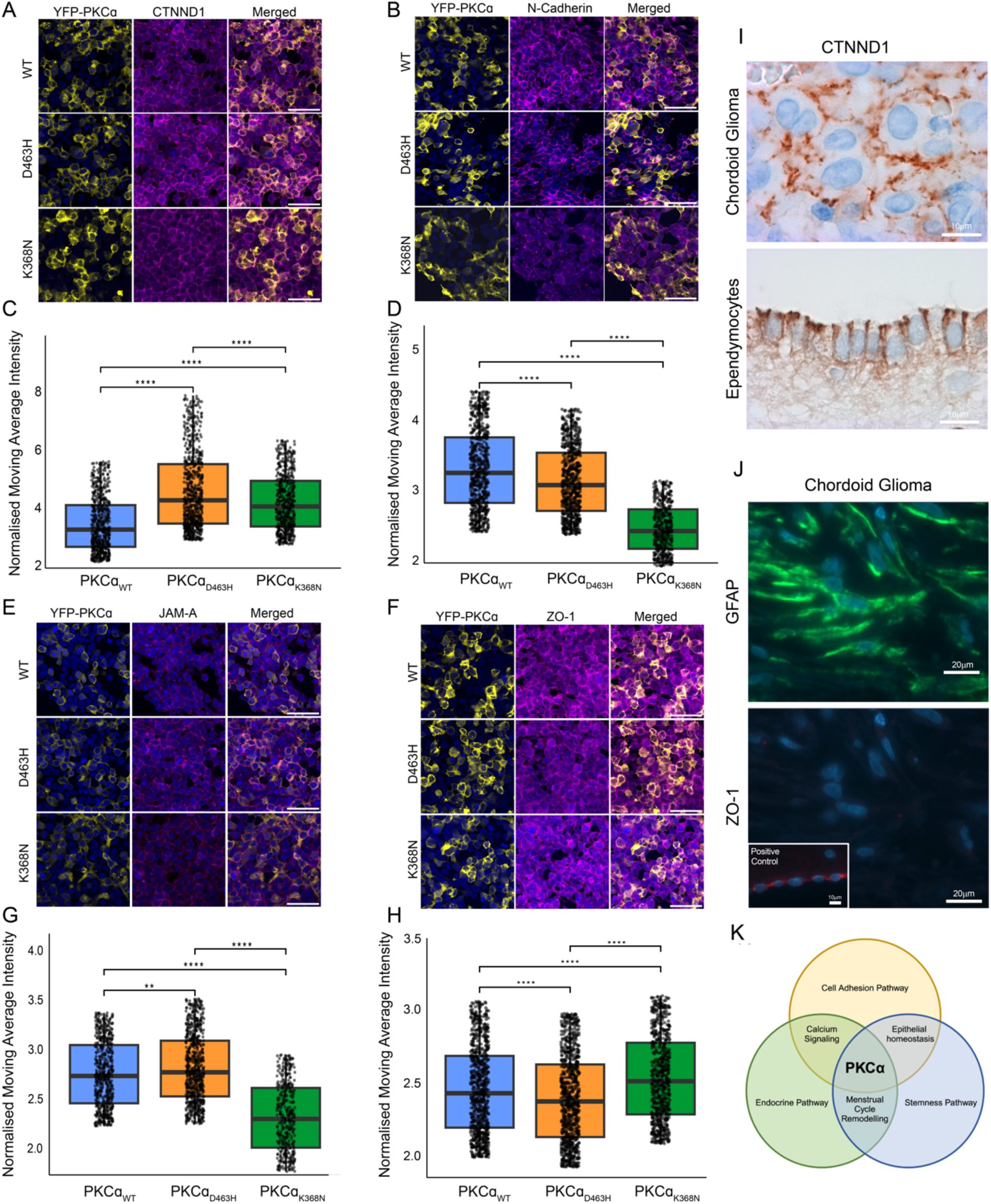
PKCα_D463H_ results in changes of localization of junction proteins at cell-cell interactions. **A-F)** Immunofluorescent staining and quantification of junctional proteins in HEK293T cells expressing YFP-PKCα_WT_, YFP-PKCα_D463H_, and YFP-PKCα_K368N._ Cells were transfected for 24 h with YFP-PKCα constructs indicated before fixing and immunostaining for **A)** CTNND1, **B)** N-Cadherin, **E)** JAM-A, **F)** ZO-1, scale bar 50 µm. Quantification of junctional protein staining intensity along the cell membranes between two YFP positive cells was performed according to custom analysis pipeline described in Figure S5 for **C)** CTNND1, **D)** N-Cadherin, **G)** JAM-A, **H)** ZO-1. (n= 1159-1620 junctions measured per condition per marker, Significance determined by Wilcoxon test for pairwise comparison, * p ≤ 0.05, ** p ≤ 0.01, *** p ≤ 0.001, **** p ≤ 0.0001). **I)** Immunohistochemical staining for CTNND1 in ChG tumor tissue (top) and ependymocytes from the hippocampus (bottom). **J)** Immunohistochemical staining of ChG tumor tissue for GFAP (top) ZO-1 (bottom). Inset – Immunohistochemical staining of ZO-1 in ependymocytes from the hippocampus. **K)** Model of how mutation in PKCα disrupts the tanycyte specific intersection between stemness, cell adhesion and endocrine function, therefore leading to chordoid glioma development.

We also observed a decrease in ZO-1 (Figure 6F, 6H), the primary marker of tight junctions, suggesting that despite JAM-A increasing at the cell membrane, the tight junction complex is lost in cells expressing YFP-PKCα_D463H_. Additionally, no ZO-1 was detected in tumor tissue (GFAP positive cells), in contrast to ependymocytes, which have tight junctions^11^ (Figure 6J). Across all measured proteins, overexpression of kinase-dead YFP-PKCα_K368N_ consistently resulted in significantly different results to YFP-PKCα_D463H_ (Figure 6A-H). These data further suggest that the effects of PKCα_D463H_ on cellular pathways are not simply due to the loss of kinase activity, but rather to a modified interactome and dominant-negative impact on PKC signaling, resulting in a significant pathway rewiring.

## Discussion

We show here that the PKCα_D463H_ mutation, the disease-defining driver mutation of chordoid glioma, rewires cell signaling pathways by a unique mechanism resulting from the specific replacement of Asp463 in the Y/HRD motif with His. This mutation not only results in a catalytically inactive kinase but also changes the stability and conformation of the protein. Our model predicts the mutation alters the dynamics of the substrate-binding surface on the I-helix to trap known substrates and aberrant binding partners in addition to binding WT enzyme. This sequestration results not only in dominant-negative effects on PKC pathway activity, but also in dysregulation of various signaling pathways, particularly those regulating cell junctions. Our data support a model in which by impairing cell polarity and adhesion, PKCα_D463H_ drives the transformation of β2-tanycytes, specialized glial barrier cells of the CVO, into ChG through their specific barrier properties.

It has been previously proposed that inhibition of PKCα could be beneficial in ChG because the D463H mutation could have catalytic activity^2^. Our data establish that the driver mutation is catalytically inactivating and dominant-negative, suppressing the activity of WT enzyme^2^. This is particularly relevant due to patients retaining one copy of WT PKCα. Thus, inhibition of PKC is inappropriate in this context and would likely be detrimental as a therapy. The functional consequences of this mutation support the prevailing tumor-suppressive role of PKC isozymes^5^ and underscore the need for therapies that restore PKC function. In this case, targeting the binding surface on the I-helix is an attractive strategy to prevent aberrant binding to unnatural binding partners. Targeting this neo protein-protein interface represents a precision oncology strategy that can neutralize the pathological effect of the mutant protein, restoring the function of WT PKC^59^.

Our study genetically confirms the cell of origin of ChG to be tanycytes, and more specifically β2-tanycytes. These specialized cells form a unilayered polarized pseudo-epithelium linked by adherens junctions on their lateral side and tight junctions on the latero-apical side of the plasma membrane^9,10^. Our coIP-MS, phosphoproteomic, and genetic data all point towards aberrant interaction, phosphorylation, and expression of proteins involved in tight junctions driven by PKCα_D463H_ (Figure 4 and 5). Loss of tight junction integrity in such a highly specialized cell could initiate tumorigenesis through impaired homeostasis and remodeling of the pseudo-epithelial layer. Through scaffolding proteins (e.g. ZO-1) and the actin cytoskeleton, tight junctions are involved in the maintenance of epithelial barrier integrity, cell polarity, and cell adhesion-dependent signaling^60^. Key tight junction proteins altered in ChG are linked with regulation of cell proliferation^61^. PKCα_D463H_ also modifies adherens junctions, whose alterations between cells is another hallmark of tumorigenesis. The main driver of such alterations has been proposed to be the loss of interaction with CTNND1 and subsequent loss of phosphorylation on Ser879, a reported PKCα site^52^. The phosphorylation of this site by PKCα has been described as a molecular switch to disassemble adherens junctions. Results from our localization studies in HEK293T cells, coupled with ChG IHC (Figure 6), support a mechanism in which loss of phosphorylation on CTNND1 stabilizes the adherens junctions and is concordant with ultrastructural studies^62^. Maintenance of adherens junctions in ChG is in line with the well-circumscribed appearance of this tumor.

PKCα_D463H_ appears to be involved in the dysregulation of GTPase signaling, as multiple proteins were observed to have altered interaction with PKCα_D463H_ alongside changes in phosphorylation and genetic expression (Figure 4 and 5). The altered proteins largely converge on the Rho GTPases Rac1 and CDC42, pointing to an alteration of the Rac1 GTPase signaling pathway, which has been associated with both junctional stability as well as focal adhesions, with a specific link to CTNND1^63–65^. In the ChG snRNAseq analysis, we see a lower enrichment for GO terms linked to focal adhesions and GTPase signaling (Figure 5E). Our proximity biotinylation analysis identified an increased interaction of PKCα_D463H_ with components of both pathways alongside microtubule assembly. Further evidence of the disruption of GTPase signaling is the loss of protein interaction of PKCα_D463H_ with with G-protein coupled receptor kinase interacting protein 1 (GIT1) alongside its partner GIT2 (Figure 4 and Table S3). The GIT1/2 complex is an important scaffold protein, controlling the spatial activation of several signaling pathways^66^, including Rac1 by recruiting it to focal adhesion complexes^67,68^. The interaction of PKCα with the GIT complex has not been previously described; however, our coIP-MS data shows a robust interaction with PKCα_WT_ which is lost with PKCα_D463H_. The dynamics of this interaction *in cellulo* are yet to be explored, particularly in the context of interactions between the ChG tumor cells and the basal membrane in the third ventricle. Taken altogether, this mutation’s impact on cell-cell junctions and cellular structure, linked with alterations of Rho GTPase signaling pathway, helps explain its role in ChG and its absence in other cancers.

The role of PKCα in tanycytes has received little attention to date. Barrier, endocrine, and homeostatic functions of tanycytes rely on the detection of and response to extracellular signals through adapting their own functional state or communicating to other cells^59^. Cell fate mapping studies have concluded that β2-tanycytes have a neuro-gliogenic capacity^57,69,70^. Studies have shown that Wnt/β-Catenin signaling inhibits the proliferation of neural progenitors, specifically hypothalamic radial glial cells^71,72^. Due to the stemness of β2-tanycytes, there could be a correlation between the downregulation of the Wnt/β-Catenin pathway, as seen in our sequencing data (Figure 5E), and the increase in proliferation of β2-tanycytes; however, this link needs to be explored further.

Tanycytes are essential neuro-endocrine cells^73,74^ that form the barrier between the CVO and surrounding brain tissue. This barrier creates an endocrine-like axis that allows for a crosstalk within the central nervous system through detection of hormones and integration of metabolic signals^75^. Plasticity is essential in endocrine tissues to allow for the modulation of physiological responses to external environmental conditions^76^. This may also make these cells more susceptible to dysregulation and mutation-induced transformation. A highly penetrant PKCα D294G mutation has been identified in tumors of endocrine tissues, specifically pituitary and thyroid tumors^77^, also causing a loss-of-function, dominant-negative effect. Given the involvement of PKCα in endocrine signaling^78^, these two highly penetrant mutations place the loss-of-function of PKCα as a key driver of tumorigenesis in endocrine tumors. PKCα has a role in the endocrine specific intersection of multiple pathways such as cellular plasticity, calcium signaling, and stemness pathways (Figure 6K). Additionally, ChG occurs with a 2:1 female predominance. Monthly remodeling of tanycytes from the estrous cycle^79,80^ may lead to the increased incidence in female patients. We hypothesize that the higher female incidence is linked to the increased plasticity of tanycytes in females. With females twice as likely to be diagnosed with ChG as males, further work should be done to understand the higher female disease burden of this tumor.

Collectively, our results reveal that PKCα_D463H_ impairs the homeostasis of the β2-tanycytes in the CVO of the lamina terminalis via a dominant-negative effect on WT PKCα signaling. We propose a model of neomorphic function leading to an abnormal network of protein-protein interactions, and dysregulation of tight junctions and adherens junctions (Figure 7). This cell junction dysregulation results in the loss of the unilayered pseudo-epithelial architecture and an abnormal, slow, and well-circumscribed growth of ChG.

**Figure 7.**
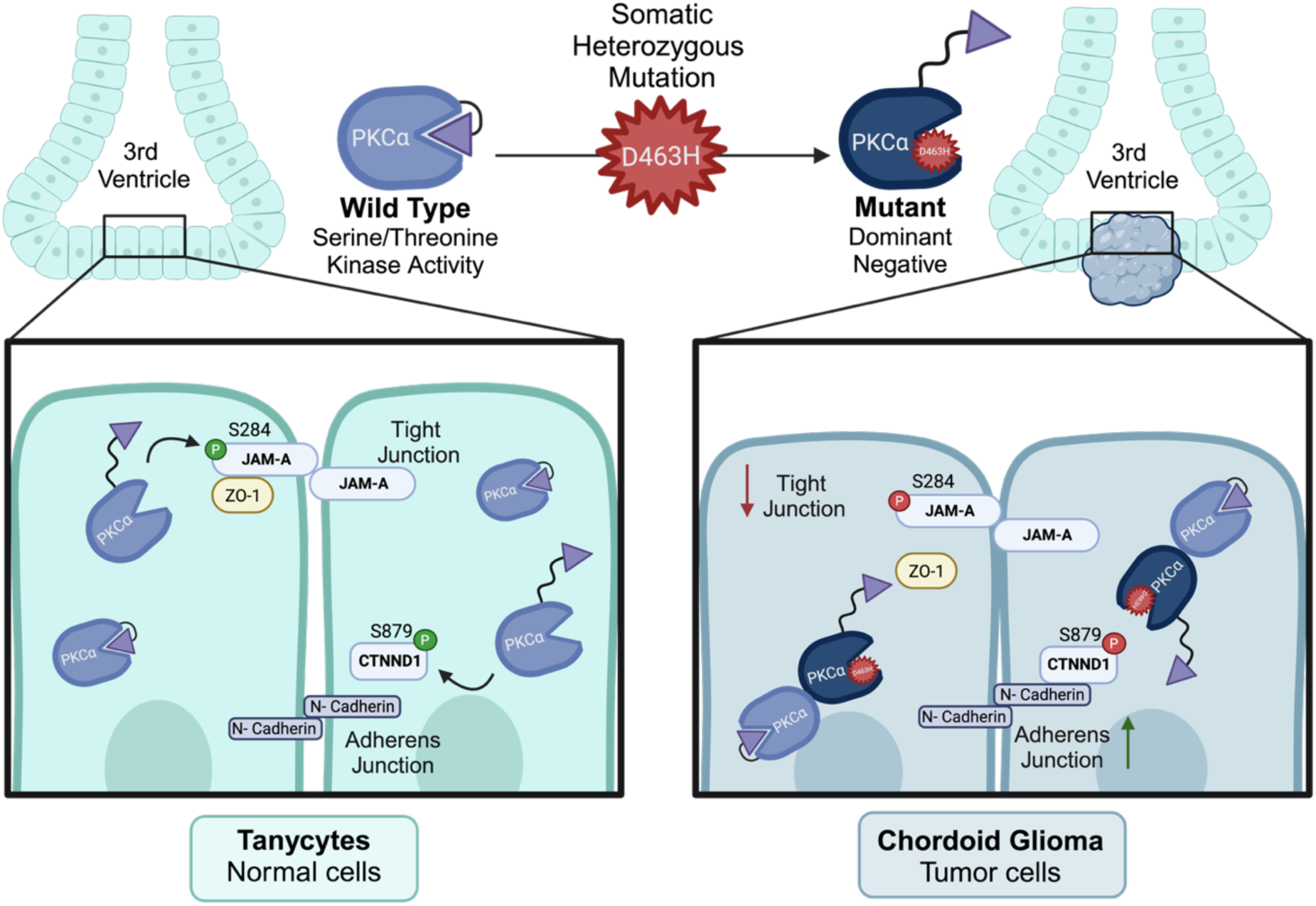
Model for the development of Chordoid Glioma from the somatic heterozygous mutation PKCα_D463H_ in Tanycyte cells of the 3^rd^ Ventricle. The PKCα_D463H_ mutation has an effect in β2-tanycytes on cellular signaling and junctional integrity, leading to ChG. This mutation rewires cellular pathways, disrupting cell-cell junctional stability, and ultimately initiating tumor formation. Here we show loss of phosphorylation at CTNND1 S879 appears to result in a stabilisation of Adherens junctions and loss of phosphorylation of JAM-A at S284 affects the localisation of ZO-1 at cell-cell interactions, likely resulting in the loss of tight junctions.

This study addresses the challenge of how a kinase mutation that does not enhance catalytic activity can still drive cancer development, through the rare example of a fully penetrant point mutation in PKCα that characterizes a low-grade glioma. This study adds to the growing literature that consolidates the role of PKCα as a tumor suppressor, whose loss-of-function ultimately leads to tumorigenesis, as well as a molecular mechanism for non-enzymatic signaling by a kinase-dead PKC mutant. This work strongly supports the cell of origin of ChG as specialized ependymal β2-tanycyte cells. The D463H mutation appears to disrupt the delicately balanced role of PKCα at the tanycyte-specific intersection of stemness, cell-adhesion, and endocrine pathways. We propose a novel precision-medicine-driven therapeutic strategy for a rare and often fatal tumor with currently no treatment options available.

### Limitations of Study

Multiple experiments in this study were carried out in HEK293T cells overexpressing PKCα. This expression system may lead to some results that are the result of overexpression of a mutant protein and not the mutation itself. However, results from the phosphoproteomic and interactomic analysis were confirmed in the snRNAseq data from ChG tumor samples. These consistencies validate our findings from HEK293T cells. There is currently no model system for ChG biology and, given the effect of PKCα_D463H_ on cell-cell junctions, an *in vivo* 3D model will be most useful to further explore the effects on tanycytes, especially as we predict there to be links with the neuroendocrine function of these cells that can only be explored in a whole-organism context.

---

**Supplementary Figure 1.**
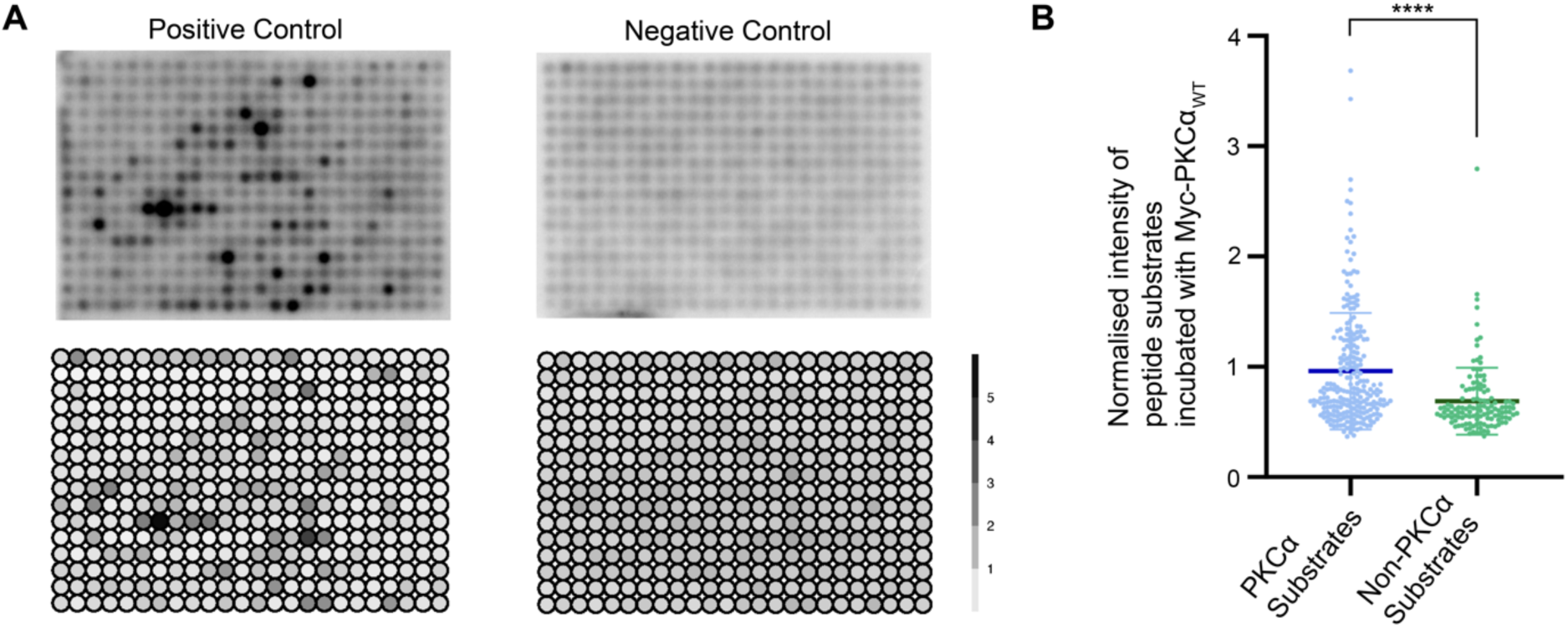
Related to figure 1. **A)** As in Figure 1B. Positive control is peptide array incubated with commercially produced recombinant human PKCα protein (Active), negative control is peptide array incubated with elution of Myc beads incubated with untransfected HEK293T cell lysates. Bottom panel shows the quantifications of n=3 averaged and normalized to negative control. **B)** Intensities from Figure 1B-C, stratified according to PKCα substrates and non-PKCα substrates, to control for specificity of phosphorylation. (**** p < 0.0001, two-tailed Mann-Whitney test)

**Supplementary Figure 2.**
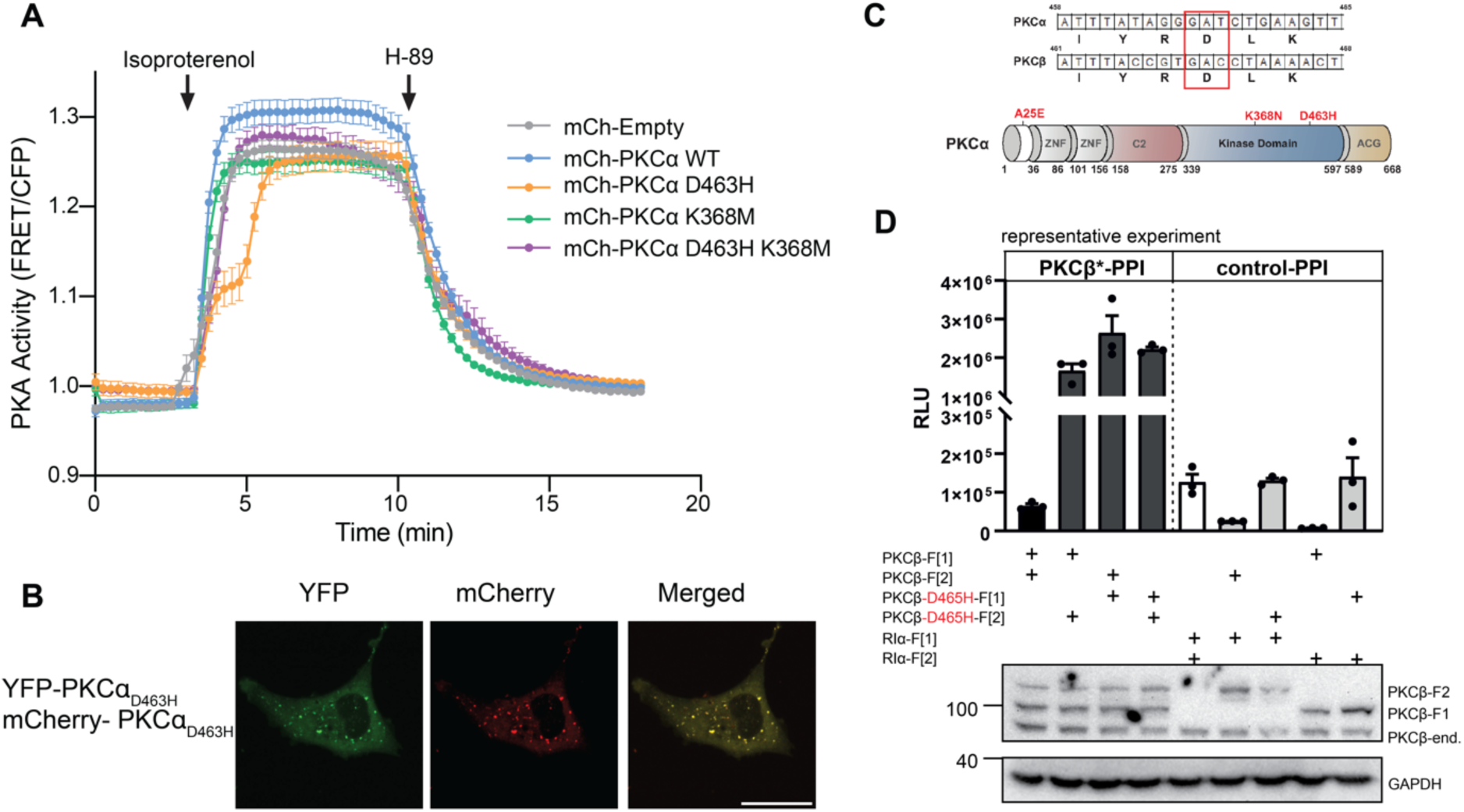
Related to figure 2. **A)** COS7 cells were transfected with AKAR PKA activity reporter and indicated mCh-tagged PKCα constructs. 24 h post-transfection, cells were treated sequentially with Isoproterenol (100 µM), and H-89 (10 µM). Data show FRET ratio changes normalized to endpoint, n=3 experiments and N≥30 individual cells, curve shows mean +/- SEM for each condition. **B)** COS7 cells grown on imaging slides were transfected with YFP-PKCα_D463H_ and mCherry-PKCα_D463H_. Cells were imaged by confocal microscopy 24 h after transfection. Scale bar: 30 µm. **C)** Upper: Alignment of partial PKCα, β sequences. Lower: Domain organization of PKCα, β; tested point mutations are highlighted in red.**D)** A representative experiment is shown of PPI analyses of PKCβ and PKA-R1α RLuc –PCA constructs. Data from one representative experiment is shown; data from technical replicates. Data are presented as relative light units (RLU).

**Supplementary Figure 3.**
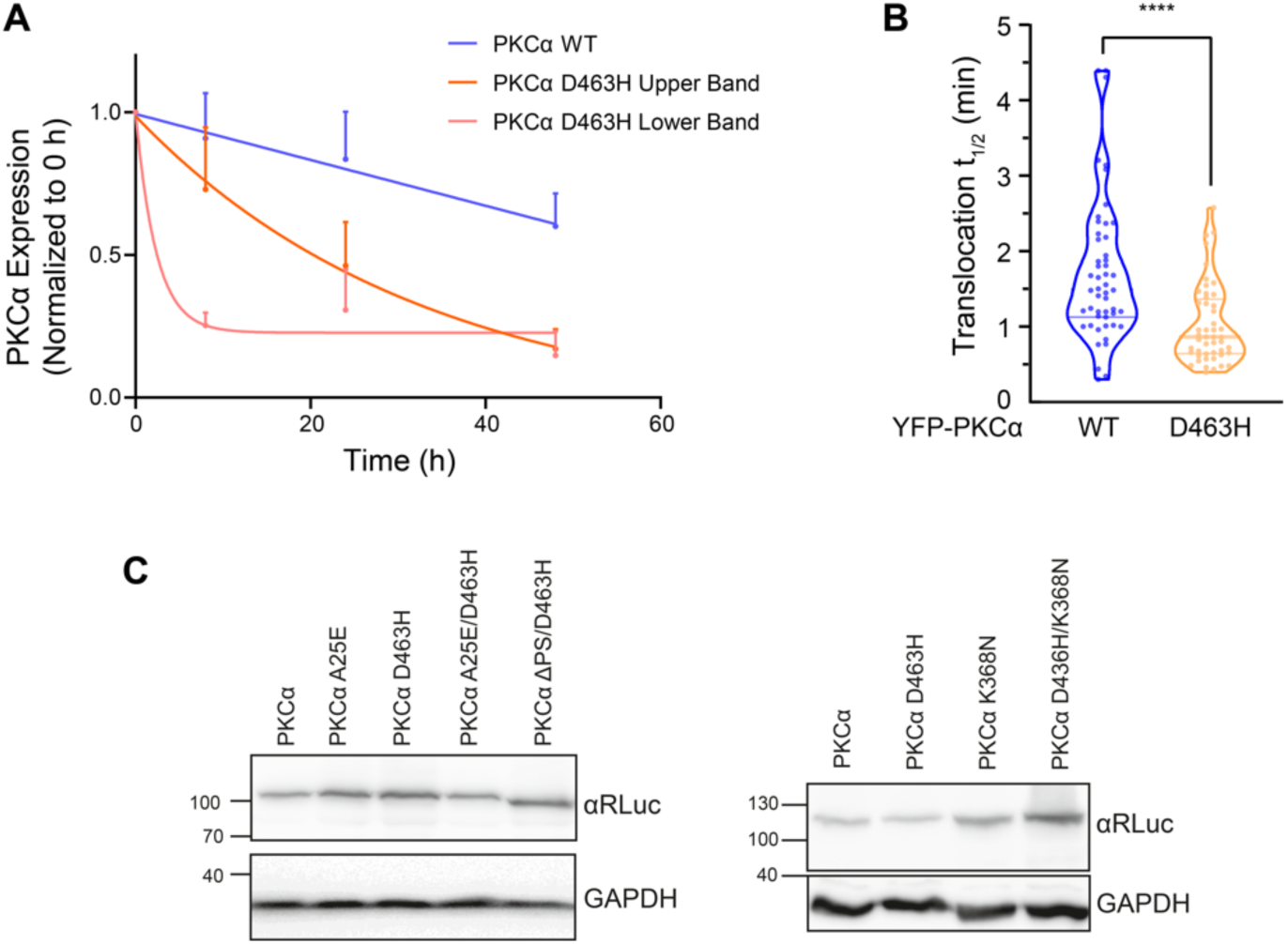
Related to figure 3. **A)** Quantification of Myc-PKCα_WT_ and Myc-PKCα_D463H_ upper and lower band protein levels from Figure 3C. Myc western blot intensity normalized to loading control and 0 h timepoint. Data were fit to first order decay, curve represents mean +/- SEM, n=3. **B)** Data for each cell from Figure 3E were fit to a non-linear regression using a one-phase association equation to calculated half-time of translocation. (**** p < 0.0001, two-tailed Mann-Whitney test). **C)** Western blots for KinCon reporter expression levels from a representative experiment are shown.

**Supplementary Figure 4.**
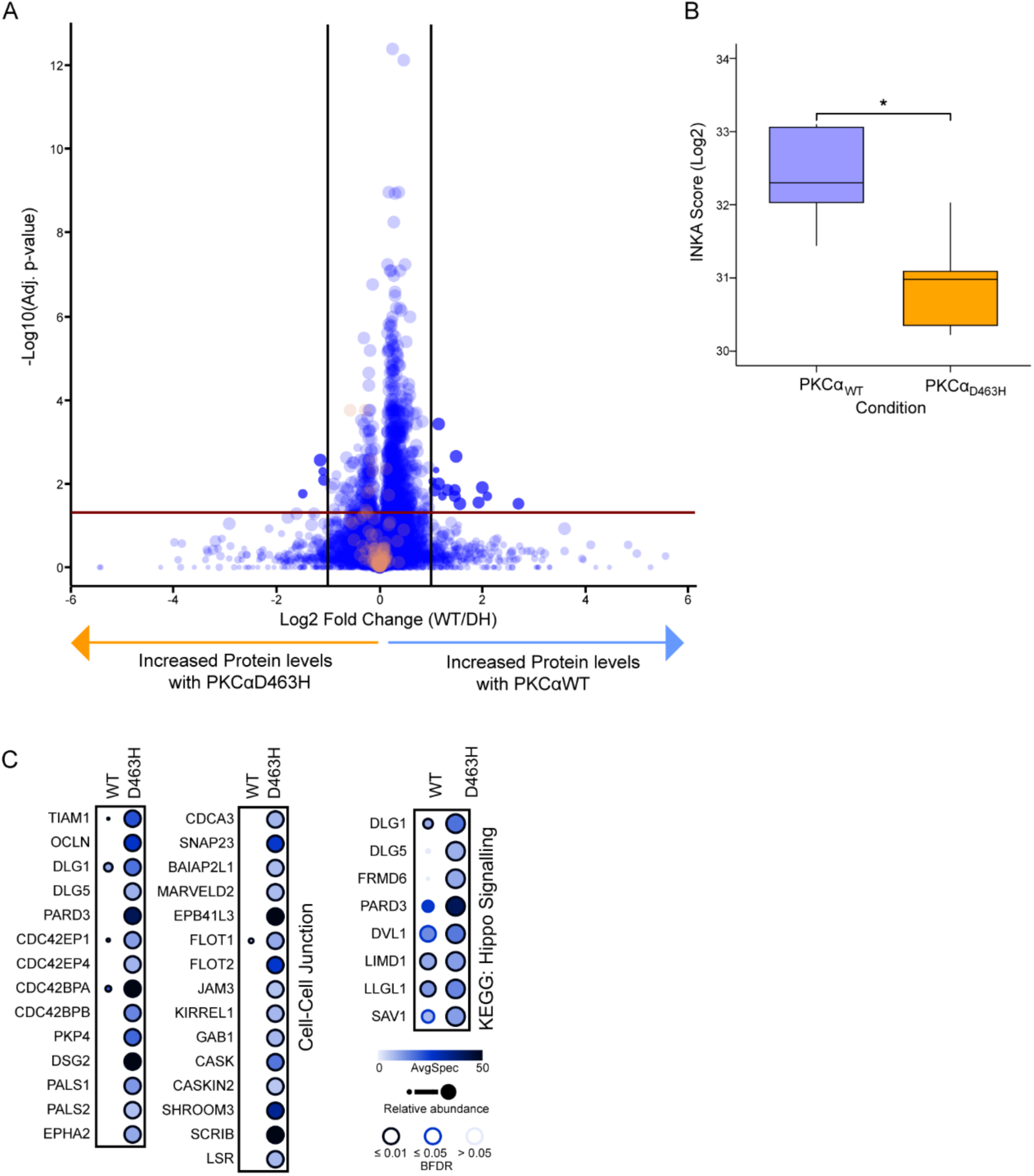
Related to figure 4. **A)** Proteome of HEK293T cells overexpressing PKCα_WT_ and PKCα_D463H_. n=5 samples per condition. Significance threshold fold change (FC)>1.5, p<0.05. Proteins with significantly altered phosphosites are highlighted in orange on the plot. **B)** INKA analysis score of significantly altered phosphosites showing the activity of PKCα in HEK293T cells overexpressing PKCα_WT_ or PKCα_D463H_. Tukey boxplots of scores shown (* p < 0.05, determined by ANOVA. **C)** Representative dot plots of proteins under enriched GO Terms found to preferentially interact with PKCα_D463H_. Dot size represents change in spectral intensity, color represents number of spectra identified, p<0.05 represented by black outline. n=2 per condition.

**Supplementary Figure 5.**
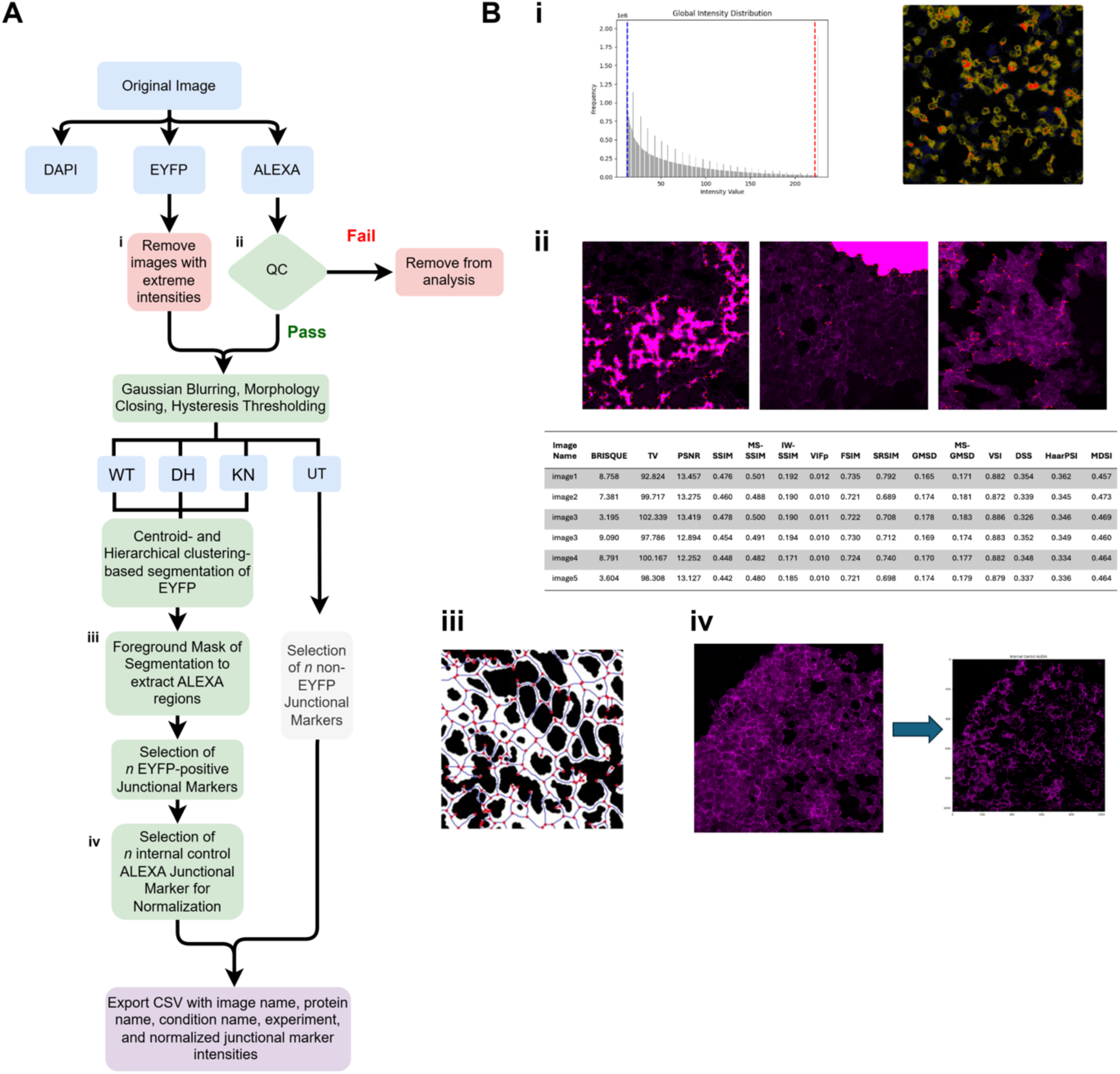
An automation method for the extraction and analyses of junctional markers in the presence of PKCα_D463H_ mutation. Related to figure 6. **A)** Overview of the automated pipeline for extracting EYFP-positive junctional markers, calibrated against the inherent internal control of each image. The pipeline involves the division of LIF (Leica Image File) formatted confocal images into their respective staining components (i.e., DAPI, EYFP, and ALEXA647 referred to as ALEXA). Image quality checks were run for the ALEXA channel using machine learning methods such as BRISQUE. Low- and high-intensity regions were excluded from EYFP images. Images then underwent pre-processing (e.g., Gaussian blurring) before EYFP regions were segmented using centroid- and hierarchical clustering-based segmentation. The EYFP segmented regions act as maps, which help with the extraction of EYFP-positive junctional markers from the ALEXA channel. Extracted intensities were normalized using the non-EYFP junctional markers within each image (given image intensities and qualities could vary over different experiments and replicates). Finally, data were automatically extracted in the form of a CSV file, ready for statistical analyses. **B)** Representative images of pipeline stages highlighted in A) **i)** The range of intensities in EYFP images were checked to ensure extreme low- and high-level intensities were removed. Low intensities may not lead to informative junctional markers, whereas highly intense regions could reflect anomalies not related to the mutation, such as cellular death. The range of the intensities were extracted on a per image basis (i.e., low-intensity regions were represented by blue and high-intensity regions represented as red), as well as the overall average lower- and upper-intensities for an experiment. EYFP regions that were either in the extreme low or high ranges were excluded from the image (i.e., blacked out), with the remaining EYFPs in the image evaluated for further analysis. **ii)** Quality assessment was run for the ALEXA channel using several statistical techniques using qualitative and quantitative measures. Assessments included natural scene statistics and machine learning-based methods, such as BRISQUE (Blind/Referenceless Image Spatial Quality Evaluator), as well as several reference-based methods, such as peak-to-signal noise ratio (PSNR) and structural similarity index measure (SSIM). Images that were outside of the expected quality thresholds were removed from further consideration. Here show examples of images which fail quality control and their respective BRISQUE scores in the table. **iii)** Extraction of the intensity of junctional markers as relative to the presence of EYFP-positive cells. Once EYFP-positive cells were selected based on pre-defined criteria (confluent and adjacent EYFP regions) and segmented, they were used as masks to then extract the equivalent junctional markers from the ALEXA-stained images. ALEXA images were converted to edges and vertices, and edges that fell within the EYFP regions segmented were extracted as a junctional marker intensity. **iv)** Normalization of intensity values against internal image controls (i.e., non-EYFP junctional markers within the same image).

## Materials and Methods

### Cell Culture and Transfection

HEK293T and COS7 cell lines were cultured in DMEM Low Glucose (Thermo Fisher Scientific Ref. 11880028), supplemented with fetal bovine serum at 10% (Thermo Fisher Scientific Ref. 10082147) and Penicillin Streptomycin at 1% (Thermo Fisher Scientific Ref. 15140122), and kept at 37 °C. Cells were transfected using Lipofectamine 3000 transfection reagent (ThermoFisher Scientific Ref. 18324012) according to manufacturer’s instructions. Sf9 cells were grown in Sf-900 II SFM media (Thermo Fisher Scientific Ref. 10902096) in shaking cultures at 27 °C.

### Plasmid Details and Mutagenesis

Full length human PKCα cDNA was inserted into mCherry, HA, YFP, and FLAG-MiniTurbo pcDNA3 vectors by Gateway Cloning (Thermo Fisher Scientific). Full length human PKCα-FLAG WT (#10805) and K368N (#10860) were purchased from Addgene. CKAR2 and MyrPalm-CFP were previously described^39^. Mutagenesis was performed by Q5 site-directed mutagenesis (New England Biosciences Ref. M0492L) or Quikchange mutagenesis (Agilent) using primers indicated in Key Resources Table.

### Protein Purification

Human GST-PKCα protein was expressed and purified from insect cells using the Bac-to-Bac expression system (Invitrogen). GST-PKCα protein was batch-purified using glutathione Sepharose beads as follows: Sf9 insect cells expressing GST-tagged protein were rinsed with PBS and lysed in 50 mM HEPES pH 7.5, 1 mM EDTA, 100 mM NaCl, 0.1% Triton X-100, 100 µM PMSF, 1 mM DTT, 2 mM benzamidine, 50 µg/mL leupeptin, and 1 µM microcystin. GST-PKCα protein was purified by incubation with GST-bind resin (Millipore Sigma Ref. 70541-5) for 30 minutes at 4 °C, washed 3 times with 50 mM HEPES pH 7.5, 1 mM EDTA, 100 mM NaCl, and 1 mM DTT, and eluted from beads with 10 mM Glutathione. Purified protein was exchanged into 20 mM HEPES, pH 7.5, 1 mM EDTA, 1 mM EGTA, and 1 mM DTT using an Amicon® Ultra-15 Centrifugal Filter Unit (Millipore Sigma Ref. UFC900308). An equal volume of glycerol was then added to a final concentration of 50% glycerol for storage at −20 °C. Protein concentration was determined against BSA standards on an SDS/PAGE gel stained with Coomassie Brilliant Blue stain.

Human Myc-PKCα was expressed and purified from HEK293T cells by transient transfection. Cells were rinsed with TBS (pH 7) supplemented with 0.1 mM Sodium orthovadanate (Thermo Fisher Scientific Ref. J60191) and lysed using M-Per mammalian Protein Extraction reagent (Thermo Fisher Scientific Ref. 78503) supplemented with Halt™ Protease Inhibitor Cocktail (Thermo Fisher Scientific Ref. 78429). Cells were lysed mechanically on ice before centrifuging at 16,000xg at 4 °C for 20 minutes. Myc-PKCα protein was purified by incubation with Pierce Anti-C-Myc-Agarose slurry (Thermo Fisher Scientific Ref. 20169) in the presence of 25% glycerol overnight at 4 °C. Beads were washed with TBS with 0.05% Tween-20 (TBS/T) five times and protein was eluted through incubation with Pierce C-Myc Peptide (Thermo Fisher Scientific Ref. 20170) at 0.5 mg/mL on a rotator for 4 hours at 4 °C. Supernatant was collected and quantified using the Qubit protein assay kit (Thermo Fisher Scientific Ref. Q33211) according to manufacturer’s instructions.

### Kinase Assays

The activity of purified GST-PKCα toward a peptide substrate (Ac-FKKSFKL-NH2) was assayed by incorporation of [32P]-phosphate from [γ-32P]-ATP into saturating amounts of substrate, measured by scintillation. 2.4 nM GST-PKCα was incubated at 30 °C in presence of 20 mM HEPES pH 7.4, 1 mM DTT, 200 µM free Ca^2+^ (as 700 µM CaCl_2_), 500 µM EGTA, 100 µM ATP (50 µCi/mL), 100 µg/mL substrate, 5 mM MgCl_2_, 0.06 mg/mL BSA, Triton X-100 (0.1% w/v), and mixed micelles containing 15 mol % PS and 5 mol % DAG in 50 mM HEPES pH 7.5 and 1 mM DTT. In non-activating conditions, CaCl_2_ was replaced with 500 µM EGTA and lipids were replaced with 20 mM HEPES. The concentration of free Ca^2+^ was calculated using a program that considers pH, Ca^2+^, Mg^2+^, EGTA, and ATP concentrations (https://somapp.ucdmc.ucdavis.edu/pharmacology/bers/maxchelator/CaMgATPEGTA-NIST.htm; Ca.Mg.ATP.EGTA program). After 5 minutes, the reaction was stopped with 100 mM EDTA pH 8.0 and 100 mM non-radiolabeled ATP. A portion of the assay mixture was spotted onto P81 phosphocellulose paper (Whatman), washed 4 times with 0.4% orthophosphoric acid, and [32P]-phosphate incorporation measured by scintillation counting.

The kinase assay peptide array was designed using known substrate peptides of PKCα and other functionally relevant substrates found on PhosphositePlus. All experimental peptides were 15 amino acids in length with a Serine or Threonine at center position, alongside 15 scramble negative controls with other amino acids at the center position, as well as the myc peptide and flag peptide. Glass peptide array slides containing 384 peptides (Table S1) were manufactured by Intavis. Slides were incubated for 4 hours with blocking buffer containing 20 mM HEPES pH 7.4, 100 mM NaCl, 5 mM MgCl_2_, 1 mM DTT, 0.2 mg/mL BSA, and 20 mM β-glycerol phosphate (Thermo Fisher Scientific Ref. J62121) at 4 °C with agitation. Slides were then washed twice with 1x Kinase Buffer (Cell Signaling Technology Ref. 9802). For the kinase assay reaction, slides were incubated for 1 hour with agitation in a humid chamber at 30 °C with 20 ng/µL Myc-PKCα in reaction buffer (1X PKC lipid activator (SignalChem Ref. L51-39), 2 µM Phorbol 12-myristate 13-acetate (PMA) (Merck Ref. P1585), 50 µM Ultra-Pure ATP (Promega Ref. V703A), 1x Kinase buffer (Cell Signaling Technology), and 90 µCi [32P] ATP (Revvity Ref. BLU002Z250UC). Purified commercial PKCα (SignalChem P61-18G-10) was used as a positive control. The reaction was stopped by washing the slide five times with 0.1 M phosphoric acid. Slides were washed five times with de-ionized water, once with methanol, and air-dried. [32P]-phosphate incorporation measured by phosphor screen on a BioRad Personal Molecular Imager FX. Analysis of spot intensities was performed using Gilles Carpentier ImageJ plugin Protein Array Analyzer for ImageJ.

### Cell Lysis and Western Blot

Cells were washed with Dulbecco’s phosphate-buffered saline (DPBS) (Corning Ref. 21-031-CV) and lysed in Phosphate Lysis Buffer pH 7.4 containing 50 mM sodium phosphate (38 mM sodium phosphate dibasic, 12 mM sodium phosphate monobasic), 1 mM sodium pyrophosphate, 20 mM sodium fluoride, 2 mM EDTA, and 1% Triton X-100. Lysis buffer was supplemented with 1 mM PMSF, 50 µg/mL leupeptin, 1 mM Na_3_VO_4_, 2 mM benzamidine, 1 µM microcystin, and 1 mM DTT added immediately prior to lysis. Whole-cell lysates were briefly sonicated prior to quantification by Bradford Assay. Samples were boiled in sample buffer containing 250 mM Tris HCl, 8% (w/v) SDS, 40% (v/v) glycerol, 80 µg/mL bromophenol blue, and 2.86 M β-mercaptoethanol for 5 minutes at 95 °C. 10 µg protein per sample was analyzed by SDS–PAGE using 6% acrylamide and transferred to Nitrocellulose membranes for 80 minutes at 80V in transfer buffer (200 mM Glycine, 25 mM Tris Base, 20% Methanol) at 4 °C. Membranes were blocked in 5% (w/v) BSA dissolved in PBS-T for 30 minutes at room temperature, washed with PBS-T 3 times for 5 minutes before incubation with primary antibody overnight at 4 °C with agitation. Membranes were washed with PBS-T 3 times for 5 minutes, incubated with secondary antibodies for 1 hour at room temperature with agitation, and washed in PBS-T 3 times for 5 minutes before developing with near-infrared fluorescent imaging. Membranes were imaged on an Azure Sapphire FL Biomolecular Imager. Antibodies –Goat anti-mouse, Azure700 conjugate – Azure Biosystems, AC2129 (1:10,000); Goat anti-rabbit, Azure800 conjugate – Azure Biosystems, AC2134 (1:10,000). Primary antibodies are described in key resources table.

### Co-immunoprecipitation Assay

Cells were transfected as described above with Myc-PKCα and FLAG-PKCα constructs. 48 hours post-transfection cells were lysed as described above and clarified by centrifugation for 10 minutes at 12,000xg. Lysates were incubated with Pierce Anti-c-myc-Agarose beads (Thermo Fisher Scientific Ref. 20169) at 4 °C overnight with agitation. The beads were washed five times with lysis buffer followed by western blot as described above.

### Cycloheximide assay

COS7 cells were transfected with 1 µg Myc-PKCα constructs for 24 hours, followed by 355 µM cycloheximide treatment for the indicated timepoints prior to lysis. Lysis and western blot protocol was followed as described above.

### Ubiquitination assay

COS7 cells were seeded into 10 cm plates and transfected with 2.5 µg YFP-PKCα DNA per plate. 48 hours post-transfection, cells were treated with 20 μM MG-132 for 4 hours followed by 200 nM PDBu for 1 hour. Cells were lysed as described above and clarified by centrifugation for 10 minutes at 12,000xg. Clarified lysates were quantified by Bradford Assay and 500 mg was used for IP. GFP-Trap® Agarose (Proteintech Ref. gab-20) or control agarose (Proteintech Ref. bab-20) was added and lysates were incubated at 4 °C overnight on a rocker with gentle motion. Beads were washed 4 times in phosphate lysis buffer and protein was eluted in sample buffer. Eluted protein was analyzed by western blot as described above.

### Kinase Conformation (KinCon reporter) Assay

HEK293T cells, cultured in StableCell DMEM (Sigma Aldrich Ref. D0822), were split into 24-well plates at 90,000 cells/well and transfected with the indicated plasmids using TransFectin Lipid reagent (Bio-Rad Ref. 1703352) at a total of 50 ng well. 48 hours after transfection, cells were washed once with PBS (1 mM Sodium phosphate pH 7.2; 15 mM NaCl), collected in 50 µL of PBS, and transferred to a 96-well plate (Grainer 96 F-Bottom). Bioluminescence was measured after adding 20 µL h-Coelentracine (Nanolight Technology) using the PHERAstar FSX (BMG Labtech) (Measurement start time: 0.2 seconds ; Measurement interval time: 10.00 seconds; Optic module LUM plus; Gain: 3600; Focal height 12.5 mm). Data was evaluated using the MARS Data evaluation Software (BMG Labtech).

### Luciferase PPI Assay

HEK293T cells were grown in DMEM supplemented with 10% FBS. The indicated RLuc-tagged constructs, tagged with either RLuc[F1] or RLuc[F2], were transiently over-expressed with TransFectin reagent (Bio-Rad Ref. 1703352) in a 24-well plate format. 48 hours after transfection, the cells were washed once with PBS (1 mM Sodium phosphate pH 7.2; 15 mM NaCl), resuspended in 150 µL PBS, and transferred to a 96-well plate (Grainer 96 F-Bottom). Bioluminescence was measured after addition of 20 µL h-Coelentracine (Nanolight Technology) using the PHERAstar FSX (BMG Labtech) (Measurement start time [s]: 0.2; Measurement interval time [s]: 10.00; Optic module LUM plus; Gain: 3600; Focal height [mm] 12.5). Data were evaluated using the MARS Data evaluation Software (BMG Labtech).

### Live-cell Fluorescent Microscopy

#### COS7 FRET imaging and analysis

COS7 cells were seeded into culture dishes (Corning Ref. 430165) containing glass cover slips (Fisherbrand Ref. 12545102) glued on using SYLGARD 184 Silicone Elastomer Kit (Dow Ref. 04019862) and transfected 24 hours after seeding. For CKAR2 assays, cells were co-transfected with 1 µg CKAR2 DNA^30^ and 1 µg mCherry-PKCα constructs or 0.1 µg mCherry alone. For translocation assays, cells were co-transfected with 1 µg YFP-PKCα and 0.1 µg MyrPalm-CFP^31^. For AKAR assays, cells were co-transfected with 1 µg mCherry-PKCα DNA and 1 µg AKAR3 DNA^81^. 24 hours post-transfection, cells were imaged in Hank’s Balanced Salt Solution (Corning Ref. 21-022-CV) supplemented with 1 mM CaCl_2_. Images were acquired on a Zeiss Axiovert 200 M microscope (Carl Zeiss Micro-Imaging Inc.) using an Andor iXonUltra 888 digital camera (Oxford Instruments) controlled by MetaFluor software (Molecular Devices) version 7.10.1.161. Background signal was subtracted for each wavelength from an area containing no cells. Regions of interest (ROI) for measurement were selected by tracing individual cells (nucleus excluded for CKAR2 and AKAR experiments). Images were acquired every 15 seconds, and baseline images were acquired for 3 minutes. Agonists and inhibitors were added as described in figure legends. For CKAR2 assays, FRET ratios for each cell were normalized to the Calyculin A maximum signal. For AKAR assays, FRET ratios were normalized to minimum signal following H89 treatment. For translocation assays, half-times were calculated by fitting the data to a non-linear regression using a plateau followed by one-phase association equation with X_0_ = 3 minutes. Data were normalized to the maximum FRET ratio for each cell (100%) to calculate percent translocation. Data represent mean +/- SEM for cells from at least three independent experiments.

#### Colocalization imaging

COS7 cells were seeded on 35 mm culture dishes with 20 mm glass bottom (Avantor VWR Ref. 734-2904) and transfected as previously described. 24 hours post-transfection, cells were imaged using the confocal microscope A1R-HD25 Nikon.

### Immunohistochemistry

HEK293T cells were grown on 12 mm glass coverslips (VWR Ref. 631-1577) after coating with Cultrex-Rat Collagen I Lower Viscosity (R&D Systems Ref. 3443-100-01) and rinsing with PBS (Corning). Cells were transfected as described above and fixed with 4% PFA (Electron Microscopy Sciences Ref. 15714) 24 hours post-transfection. Cells were blocked for 1 hour at room temperature with 0.3% Triton X-100 and 4% Normal Goat Serum (Sigma-Aldrich Ref. S26) in PBS. Blocked cells were washed three times with PBS then incubated overnight at 4°C in primary antibody diluted in blocking solution. Antibodies used: ZO-1 (1:200), Rabbit IgG (Thermo Fisher Scientific Ref. 40-2200); CTNND1 (1:800), Rabbit IgG (Cell Signaling Technology Ref. 59854); JAM-A (1:400), Rabbit IgG (Cell Signaling Technology Ref. E6Z7E); N-Cadherin (1:600), Rabbit IgG (Cell Signaling Technology Ref. D4R1H). Coverslips were then rinsed three times with PBS and incubated for 1 hour at room temperature with corresponding secondary antibody in blocking solution. Coverslips were washed three times with PBS and mounted onto slides using Fluoroshield with Dapi Histology mounting medium (Sigma Ref. F6057). Slides were then imaged using Confocal SP8 X White Light Laser Leica and analyzed in ImageJ.

For ChG tumor staining, 4 mm sections of formalin-fixed paraffin-embedded tumor tissue were deparaffinized and labelled by a fully automatic Ventana benchmark Ultra System (Roche, Basel, Switzerland) using a streptavidin-peroxidase complex with diaminobenzidine as the chromogen and the following primary antibody CTNND1 (1:800), Rabbit IgG, (Cell Signaling Technology Ref. 59854), GFAP (1:400), Mouse mAB (Cell signaling Technology Ref: 3670BF), ZO-1 (1:400), Rabbit IgG (Abcam Ref: ab221547).

### Molecular Modeling

Molecular dynamics (MD) simulations were performed on both the WT and D465H mutant forms of PKCβ (PDBID: 2I0E) using AMBER16. Each system underwent five replicate simulations for 10 nanoseconds. Structures were extracted every 100 picoseconds, resulting in 100 structures per replicate for analysis. The Local Spatial Pattern (LSP) alignment method was used to align all 100 structures in an all-to-all manner, producing 4950 alignments per trajectory, as previously described^37^. The results were averaged across the alignments, and this process was repeated five times for each construct. The final values, along with the standard error (SE), were calculated from these five replicate simulations.

### Phosphoproteomic, Proteomic, and Co-Immunoprecipitation Mass Spectrometry

#### Sample preparation

For co-immunoprecipitation mass-spectrometry, HEK293T were transfected with PKCα_WT_-FLAG and PKCα_D463H_-FLAG using the calcium phosphate transfection method. 48 hours post-transfection, cells were washed in PBS, dissociated from the dish by trypsin before lysis with 1X Cell Lysis Buffer (Cell Signaling Technology Ref. 9803S) consisting of 20 mM Tris-HCL (pH 7.5), 150 mM NaCl, 1 mM Na_2_EDTA, 1 mM EGTA, 1% Triton X-100, 2.5 mM sodium pyrophosphate, 1 mM β-glycerophosphate, 1 mM Na_3_VO_4_, and 1 μg/mL leupeptin) and Halt™ Protease Inhibitor Cocktail (Thermo Scientific Ref. 78438). Lysates were precleared with protein G magnetic beads (Cell Signaling Technology Ref. 70024) for 20 minutes, before incubation with DYKDDDDK Tag (9A3) Mouse mAb (Cell Signaling Technology Ref. 8146) overnight with rotation at 4 °C. The beads were washed three times with 25 mM NH_4_HCO_3_, resuspended in 100 μL of 25 mM NH_4_HCO_3_, and digested by adding 0.2 μg of trypsin/LysC (Promega) for 1 hour at 37 °C. Samples were loaded into custom-made C18 StageTips packed by stacking one AttractSPE® disk (Affinisep Ref. SPE-Disks-Bio-C18-100.47.20) and 2 mg beads (Waters Ref. 186004521) into a 200 µL micropipette tip for desalting. Peptides were eluted using a ratio of 40:60 MeCN:H_2_O + 0.1% formic acid and vacuum concentrated to dryness. Peptides were reconstituted in 10 µL of injection buffer (0.3% TFA) before liquid chromatography tandem mass spectrometry (LC-MS/MS) analysis.

For proteomic and phosphoproteomic analysis, HEK293T cells were transfected as described above and lysed in freshly prepared urea buffer (8 M urea, 200 mM ammonium bicarbonate). After sonication, lysates were quantified by BCA and 250 µg of each cell lysate was reduced with 5 mM DTT for 1 hour at 37 °C before alkylation with 10 mM iodoacetamide for 30 minutes at room temperature in the dark. Samples were then diluted in 200 mM ammonium bicarbonate to reach a final concentration of 1 M urea and proteins were digested overnight at 37 °C with trypsin/Lys-C (Promega Ref. V5071) at a ratio of 1:50. Digested peptides were acidified with formic acid to a final concentration of 5% formic acid. Samples were centrifuged at 4000 rpm and loaded onto homemade SepPak C18 Tips for desalting as described above by using three AttractSPE® disk and 20 mg beads. Peptides were eluted and 90% of the starting material was enriched using Titansphere^TM^ Phos-TiO kit centrifuge columns (GL Sciences Ref. 5010-21312, GL Sciences) as described by the manufacturer. After elution from the Spin tips, the phospho-peptides and the remaining 10% eluted peptides were vacuum concentrated to dryness and reconstituted in 0.1% formic acid prior to LC-MS/MS phosphoproteome and proteome analyses.

#### LC-MS/MS Analysis

Peptides from IP samples were separated by reverse phase liquid chromatography (LC) on an RSLCnano system (Ultimate 3000, Thermo Fisher Scientific) coupled online to an Orbitrap Fusion Tribrid mass spectrometer (Thermo Fisher Scientific). Peptides were trapped in a C18 column (75 μm inner diameter × 2 cm; nanoViper Acclaim PepMap 100, Thermo Fisher Scientific) with buffer A (2:98 MeCN:H2O in 0.1% formic acid) at a flow rate of 4.0 µL/minute over 4 minutes. Separation was performed on a 50cm x 75μm C18 column (nanoViper Acclaim PepMap^TM^ RSLC, 2 μm, 100Å, Thermo Fisher Scientific) regulated to a temperature of 55 °C with a linear gradient of 5% to 25% buffer B (100% MeCN in 0.1% formic acid) at a flow rate of 300 nL/minute over 100 minutes. MS1 data were collected in the Orbitrap (120,000 resolution; maximum injection time (IT) 50 milliseconds; AGC 4E5) and MS2 scan performed in the ion trap in rapid mode with high energy collision dissociation (HCD) fragmentation (isolation window 1.6 Da; NCE 30%, IT 100 milliseconds; AGC 2E4).

For proteome and phosphoproteome samples, LC was performed with an RSLCnano system coupled to a Q Exactive HF-X mass spectrometer (Thermo Fisher Scientific). Peptides were trapped onto the C18 column with buffer A at a flow rate of 2.5 µL/minute over 4 minutes. Separation was performed at a temperature of 50°C with a linear gradient of 2% to 30% buffer B at a flow rate of 300 nL/minute over 91 minutes for the phosphoproteome analyses and a linear gradient of 2% to 35% buffer B over 211 minutes for the proteome analyses. MS full scans were performed in the ultrahigh-field Orbitrap mass analyzer in ranges m/z 375–1500 (120,000 resolution; IT 50 milliseconds; AGC 3E6). The top 20 most intense ions were subjected to Orbitrap for further fragmentation via HCD activation (15,000 resolution; AGC 1E5; NCE 27). We selected ions with charge state from 2+ to 6+ and the dynamic exclusion to 20 seconds for the phosphoproteome and 40 seconds for the proteome analyses.

#### Data analysis

For identification, the data were searched against the *Homo Sapiens* (UP000005640) database (downloaded 01/2018) using Sequest HT through Proteome Discoverer (version 2.2). Enzyme specificity was set to trypsin and a maximum of two missed cleavage sites were allowed. Oxidized methionine, N-terminal acetylation, carbamidomethylation of cysteine were set as variable modifications. Phosphorylated serine, threonine, and tyrosine were also set as variable modifications for IP and phosphoproteome samples. Maximum allowed mass deviation was set to 10 ppm for monoisotopic precursor ions. For fragment ions, it was set respectively to 0.6 Da and 0.02 Da for the Orbitrap Fusion and the Q Exactive HF-X data. The resulting files were further processed using myProMS^82^ [https://github.com/bioinfo-pf-curie/myproms] v.3.10. False-discovery rate (FDR) was calculated using Percolator ^83^ and was set to 1% at the peptide level for the whole study. Label-free quantification was performed using peptide extracted ion chromatograms (XICs) and computed with MassChroQ^84^ v.2.2.1.

XICs from peptides shared between compared conditions (TopN matching for IP and proteome conditions and sample ratio for phosphoproteome) with no missed cleavages were used. Median and scale normalization at peptide level was applied on the total signal to correct the XICs for each biological replicate (N=5). The phosphosite localization accuracy was estimated by using PhosphoRS in myProMS. Phosphosites with a localization site probability greater than 95% were quantified at the peptide level. To estimate the significance of the change in protein abundance, a linear model (adjusted on peptides and biological replicates) was performed and p-values were adjusted using the Benjamini–Hochberg FDR procedure.

For coIP-MS data: GO term enrichment analysis was performed using a list of proteins with at least 2 distinct peptides across 3 biological replicates in the FLAG-PKCα_WT_ versus FLAG-PKCα_D463H_ condition. A 1.5-fold enrichment and an adjusted p-value ≤ 0.05 were considered significantly enriched in sample comparisons. Graphical representation shows the 1.5-fold change with 2 distinct peptides across 3 biological replicates and an adjusted p-value ≤ 0.05. Highlighted in grey are proteins that correspond to Keratinocyte related GO terms: GO:0005840; GO:0022626; GO:0030532; GO:0045095.

For Phosphoproteome data: GO term enrichment analysis was performed using a list of proteins with at least 1 distinct peptide across 3 biological replicates. A 1.5-fold enrichment in the FLAG-PKCα_WT_ vs FLAG-PKCα_D463H_ condition and an adjusted p-value ≤ 0.05 were considered significantly enriched in sample comparisons. Unique proteins were considered with at least 1 total peptide in all replicates. Graphical representation shows proteins with at least 1 distinct peptide across 3 biological replicates, a 1.5-fold enrichment in the PKCα_WT_ vs PKCα_D463H_ condition, and an adjusted p-value ≤ 0.05.

For Proteome data: Graphical representation shows proteins with at least 3 total peptides in all replicates, a 2-fold enrichment, and an adjusted p-value ≤ 0.05 were considered significantly enriched in sample comparisons. The resulting files were further processed using myProMS. GO enrichment analysis was performed by the perl module GO:Termfinder (https://metacpan.org/dist/GO-TermFinder)^85^, integrated in myProms (FDR<=1%, Benjamini & Hochberg), using the Complete Gene Ontology (v. 2023-03-06, 2023-03-06) Uniprot-GOA Human (v. 2023-02-07)^49,50^.

The data has been deposited at PRIDE, https://www.ebi.ac.uk/pride/ under the identifier : PXD056944 (Username, Password: N16DBl9mwGpI)

### INK-A Analysis

Kinase activity was inferred from phosphoproteomic data using Integrative iNferred Kinase Activity (INKA) analysis as previously described^48^. INKA scores are calculated based on four parameters: the sum of all phosphorylated peptides belonging to a kinase, the detection of the phosphorylated kinase activation domain (kinase-centric parameters), the detection of known phosphorylated substrates, and the presence of predicted phosphorylated substrates (substrate-centric parameters). The latest version of the INKA pipeline is available online at https://inkascore.org/ and is fully integrated in myProms. ANOVA was performed on Inka score between the 2 groups (FLAG-PKCα_WT_/FLAG-PKCα_D463H_).

### Proximity Biotinylation

Sample Generation and Lysis: Flp-In™ T-REx™ 293 stable cell lines (Invitrogen) expressing FLAG-MiniTurbo-empty, PKCα_WT_, or PKCα_D463H_ mutant were induced for 24 hours with 0.1 µg/mL doxycycline for protein expression. Proximity biotinylation of interacting proteins was activated by the addition of 50 µM Biotin (Sigma) for 1 hour before lysis. Cells were washed twice in ice-cold PBS (Corning) and pelleted at 500xg for 5 minutes before freezing at -80°C. Frozen cell pellets were resuspended in ice-cold modified RIPA lysis buffer (50 mM Tris-HCl pH 7.4, 150 mM NaCl, 1 mM EGTA, 0.5 mM EDTA, 1 mM MgCl_2_, 1% NP40, 0.1% SDS, 0.4% sodium deoxycholate, 1 mM PMSF , and 1x Protease Inhibitor cocktail) at a 1:4 pellet weight:volume ratio. Cells were sonicated for 15 seconds on ice, treated with 250 U TurboNuclease and 10 µg RNase, and rotated (end-over-end) at 4 °C for 15 minutes. 10% SDS was added to further increase the SDS concentration to 0.4% before rotating the cells at 4 °C for 5 minutes. Cells were centrifuged at 20,817xg for 20 minutes at 4 °C and supernatant was transfered to a 2 mL centrifuge tube.

Biotin-Streptavidin Affinity Purification: 20 μL of streptavidin beads were added to clarified lysate and incubated with rotation for 3 hours at 4 °C. Beads were pelleted 400xg for 1 minute at 4 °C, washed once with SDS-Wash buffer (25 mM Tris-HCl pH 7.4, 2% SDS), twice with RIPA-wash buffer (50 mM Tris-HCl pH 7.4, 150 mM NaCl, 1 mM EDTA, 1% NP40, 0.1% SDS, 0.4% sodium deoxycholate), once with TNNE buffer (25 mM Tris-HCl pH 7.4, 150 mM NaCl, 0.1% NP40, 1 mM EDTA), and 3 times with 50 mM ammonium bicarbonate (ABC) buffer pH 8.0.

Trypsin Digest and MS Analysis, Data Analysis: Beads were incubated at 37 °C overnight with 1 µg trypsin in 50 mM ABC buffer with agitation. An additional 0.5 µg trypsin was added for 3 hours. Supernatant was collected after centrifugation at 400xg for 2 minutes, clarified by centrifugation at 16,000xg for 10 minutes, and lyophilized by vacuum centrifugation until dry. Peptides were identified by LC-MS/MS on Bruker timsTOF Pro 2 and analysis with MSFragger 3.7, against UP000005640 human Uniprot database. Significant interactors were identified by SAINT Analysis with a Bayesian False Discovery rate <=0.01.

### snRNAseq

#### Single-nuclei preparation

Single nuclei were isolated using an in-house protocol, as previously described^86^. Briefly, frozen tissue samples were minced in a lysis buffer (10 mM Tris–HCl, 10 mM NaCl, 3 mM MgCl₂, and 0.1% Nonidet™ P-40 in nuclease-free water) and mechanically dissociated using A and B pestles, gently for 15 times each. Samples were then suspended in 2% BSA in PBS, sieved through 100 µm cell strainers (VWR, Ref. 732-2759P), and centrifuged twice for 10 minutes at 500xg. The resulting pellet was resuspended in 2% BSA in PBS. Nuclei were incubated with an Alexa Fluor® 647 anti-Nuclear Pore Complex Proteins Antibody (BioLegend Ref. Mab414), then sorted using a FACSAria™ III (BD Biosciences) with the 85 µm nozzle and the BD FACSDIVA™ software. Sorted nuclei were immediately processed on a Chromium™ Controller (10x Genomics).

#### 5’ Gene expression single nuclei library preparation

Single-nuclei suspensions (target capture of 10,000 nuclei) were loaded onto the Chromium Controller microfluidics device and processed following the Chromium Single Cell 5′ Library V2 protocol (10x Genomics, Pleasanton, CA). After the quality control of the cDNA amplification, 5’ gene expression Dual Index libraries were prepared, quality controlled, and sequenced on HighOutput flowcels of Nextseq 500 to obtain the recommended number of 20,000 reads per targeted nuclei. Between 4200 and 5640 nuclei per sample were identified (for 1260-1810 detected genes).

#### snRNAseq

snRNA processing: The Cell Ranger^8787^. Single-cell Software suite (4.0.0) was used to process the data with GRCh38-1.2.0_premrna reference genome. Filtered_feature_bc_matrix outputs were loaded into Seurat^8888^ bioconductor package v4.4.0 (R v4.1) to filter the datasets and identify cell types. Genes expressed in at least 5 nuclei and nuclei with at least 200 features were retained for further analysis. To remove likely dead or multiplet nuclei from downstream analyses, nuclei were discarded when they had less than 500 UMIs (Unique Molecular Identifiers), greater than 20,000 UMIs, or expressed over 8 % mitochondrial genes. Gene expression matrix was log normalized using the method implemented in the Seurat’s *NormalizeData* function. To cluster nuclei, we computed a Principal Components Analysis (PCA) on scaled variable genes, as determined above, using Seurat’s *RunPCA* function, and visualized it by computing a Uniform Manifold Approximation and Projection (UMAP) using Seurat’s *RunUMAP* function on the top 30 PCs. We also computed the k-nearest neighbor graph on the top 30 PCs using Seurat’s *FindNeighbors* function and in turn used Seurat’s *FindClusters* function with varying resolution values. We chose a final value of 0.3 for the resolution parameter. The *FindMarkers* function with the default parameters was used to identify differentially expressed genes in different clusters.

1. Deconvolution of RNA samples based on snRNA ChG sample clusters using cybersortX^89^: Normalized counts of ChG sample and nuclei cluster information were used as input of cybersortX to generate a signature matrix of our population clusters. The signature matrix was then used to deconvolute normalized (counts per million) counts of our 9 RNA seq samples by imputing cell fractions with absolute mode and default parameters.
2. Differential analysis cluster 1 and human astrocyte: We merged raw counts of the two datasets, do a normalization step with Seurat’s *SCTransform* function. We used the cluster labels determined previously on separated objects for differential analysis using Seurat’s *FindMarker*. Differentially expressed genes were then used as input to gProfiler for enrichment analysis.
3. Comparison of mouse tanycytes scRNA and snRNA in ChG: We used a published dataset of scRNA of murine tanycytes^56^. To be able to compare the murine scRNA with our human snRNA of ChG, we converted mouse genes names to human gene symbols using an orthologous genes database. This was achieved using BiomartRt’s^90^ *useMart* for human and mouse and *getLDS* to connect the mart objects and extract wanted information.Tanycyte signatures for each cluster was determined by using the up-regulated genes for each tanycyte suptype with Seurat’s *AddModuleScore*.

The data discussed in this publication have been deposited in NCBI’s Gene Expression Omnibus^91^ and are accessible through GEO Series accession number GSE279603 (https://www.ncbi.nlm.nih.gov/geo/query/acc.cgi?acc=GSE279603).

### Statistical Analysis

Statistical methods applied are as described in figure legends and performed in Prism version 9.4.1.

### Analysis pipeline for detection of junctional markers from EYFP-positive cells

A pipeline was developed for the extraction of intensities from the junctional markers. The pipeline includes the splitting of the images into their respective stains (i.e., DAPI, EYFP, and ALEXA). Initially, we assessed the intensity ranges of EYFP positive cells, as low intensity regions would not be assisted in the evaluation of the *PRKCA* D463H mutation, while high intensity regions would be reflective of other phenomenon, such as cellular death. The pipeline includes iteratively evaluating all EYFP-stained images, assessing the lower and upper intensities across each image individually before assessing the average across all images collectively. These ranges were then used to “black out” or remove the low and high intensity regions. In parallel to the EYFP assessment, ALEXA-stained images also underwent additional image quality assessment. Image quality assessment included a combination of no-reference, machine learning based measures (i.e., BRISQUE), as well as reference-based statistics, such as peak signal-to-noise ratio (PSNR), structural similarity index measure (SSIM), and multi-scale SSIM (MS-SSIM). Detailed summary of all metrics can be found in **Supplementary Information**. For reference-based statistics, the best quality images were selected by the author CB as the reference image. The images were assessed on a patch-level basis, with any regions falling outside of the ideal threshold for quality being marked with red dots on the image. Images that were marked as low quality both visually and quantitively were removed from further analyses. Once clean and good quality images were selected, binarized masks were created using image pre-processing methods including Gaussian blurring, morphology closing, and hysteresis thresholding. The four conditions were separated into two different parts of the pipeline. The first, which included the DH mutant, KN control, and WT, involved the segmentation of pre-defined EYFP-positive cells using centroid- and hierarchical clustering-based segmentation. The EYFP-positive cells selected included those which were confluent and adjacent to one another (to ensure the relevant junctional markers were being extracted). This segmentation led to a binarized mask of the cells of interest being generated. These masks were then used to map which junctional markers needed to be extracted from the ALEXA images. In binarized ALEXA images, images were mapped using edges and vertices, with vertices representing the corners of the cells and the edges the junctional markers. EYFP-positive celled masks that correlated with these junctional markers would be extracted. Extracted junctional markers additionally underwent a normalization process to account for potential intensity variabilities across different images within different technical replicates and experiments (i.e., intra-rater variability). This normalization involved identifying non-EYFP-impacted junctional markers from the same image in which the EYFP-positive intensities were extracted, getting the average of these non-EYFP markers using a moving average (as even individual images there can be variable in intensities, which may not be captured with the selection of a handful of randomly selected control junctional markers), and using this moving average to normalize captured EYFP-positive junctional markers by dividing its intensity with the moving average. This process was iterated through all images automatically, with the result being a singular CSV file that contained variables, such as the image name, protein, condition, experiment and replicate, location of junctional marker, the moving average for the image, and the normalized intensity of the junctional marker.

The statistical difference between the different conditions were assessed for each protein and replicated using boxplots and jitter. Boxplots were generated to visualize distributions of normalized intensity across different conditions. For pairwise comparisons, the Wilcoxon test was applied, ensuring that there were enough data points for each tested condition.

## Supplemental Information

**Figures S1–S5 and Table S1-3**

**Table S1.** List of peptides and corresponding proteins used in Peptide array, related to Figure 1

**Table S2.** List of significantly altered phosphosites, related to Figure 4

**Table S3.** List of significantly altered binding partners from coIP-MS analysis, related to Figure 4

## Acknowledgements

The authors thank the staff of ICM platforms Histomics (histology), DAC (Data Analysis Core), Quant (microscopy), and Celis (cell culture). They are grateful to Amel Dridi-Aloulou and the biobank Onconeurotek (AP-HP) for the cession of frozen tumor samples. They are grateful to the technical staff of Neuropathology lab Raymond Escourolle for the preparation of tissue section and chromogenic immunostaining. C.B. thanks Arnaud Klein and Alain Sureau from the Centre de Recherche de Myologie for using the phosphorimager and screens for the peptide array as well as Eric Noe for his assistance with radioactive safety. The LSMP thanks Patrick Poullet from the bioinformatics platform of the Institut Curie U900 for the continuous development of myProMS. The authors wish to thank Laura McGary of the Network Biology Collaborative Centre Proteomics Facility (RRID: SCR_025375) at the Lunenfeld-Tanenbaum Research Institute for proximity-dependent biotinylation sample processing, mass spectrometry, and data analysis. The facility is supported by the Canada Foundation for Innovation and the Ontario Government. Thanks to Celine Bertholle and Vaarany from the Cochin Institute CYBIO core facility for the technical help in the Single nuclei RNAseq experiments.

## Author contributions

C.B. and H.T. designed and performed the experiments, wrote the manuscript and drafted the figures. S.S. performed the PCA assays under the supervision of E.S. Q.L. analyzed the snRNAseq and bulk RNAseq data under supervision of C.B. and F.B. D.L. oversaw the Phosphoproteomic, Proteomic and coIP-MS experiments assisted by F.D. J.A. created the image analysis pipeline and quantified junctions. A.K. performed the structural analysis. J.L. performed IHC of ChG tissue. T.H.K. and T.R.B. performed experiments under the supervision of A.C.N. S.L. contributed to *in silico* analysis of Phosphoprotemic data. B.I. and M.A. performed the snRNAseq of ChG. H.A.-B. recruited tumor samples. J.-V.B. contributed to the peptide array. F.B, M.S and A.C.N conceived the project, designed the experiments, and wrote the manuscript.

## Data and material availability

Further information and requests for resources and reagents should be directed to and will be fulfilled by the lead contacts, Franck Bielle and Alexandra Newton. Plasmids generated in this study are available upon request to the lead contact. Other materials are available through commercial sources (see resource table). Data produced in this paper are available online at depository sites listed in the key resource table and materials and methods and will be made public upon publication. Code for the image analysis pipeline has been uploaded and will be made public upon publication.

## Resource Table

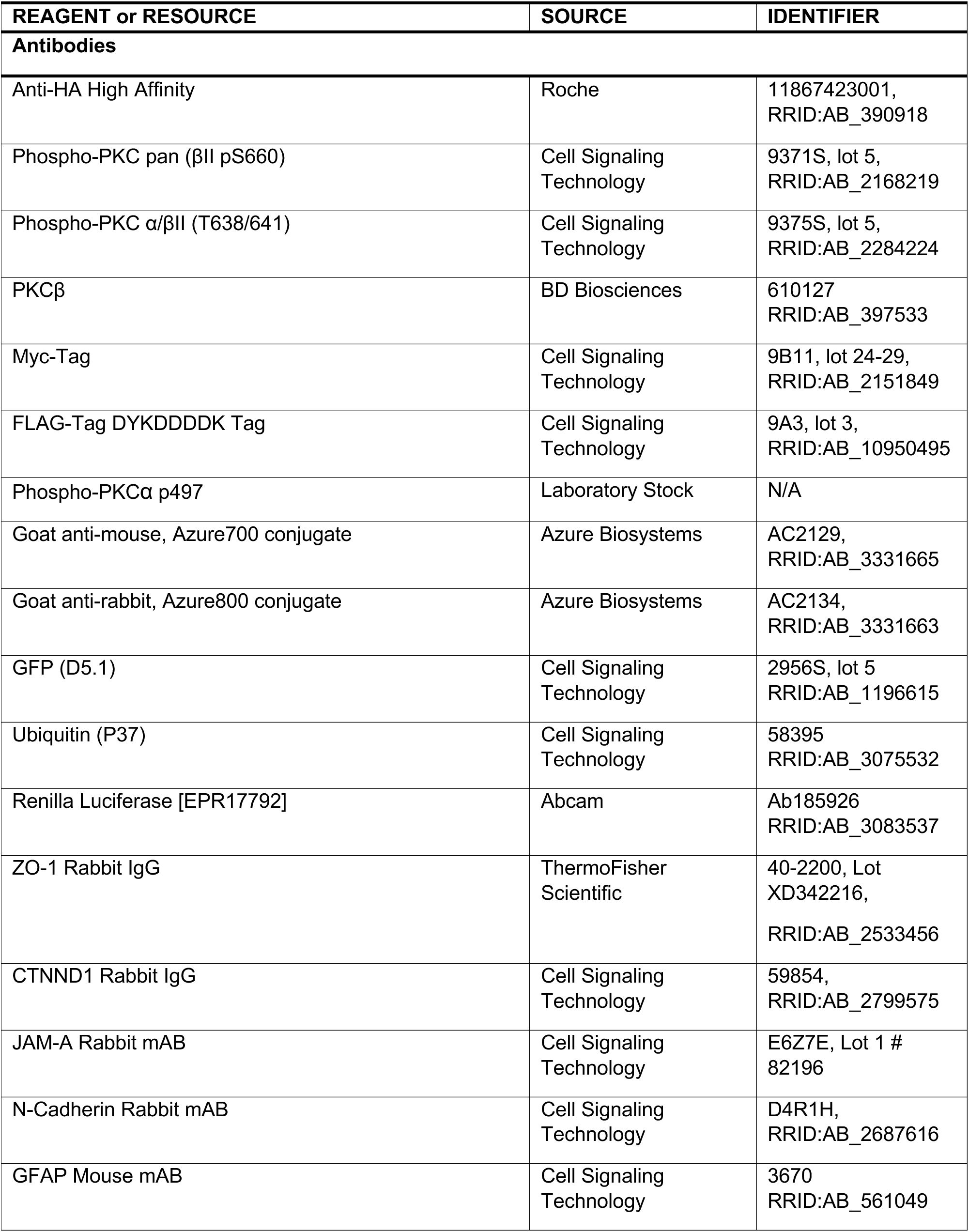
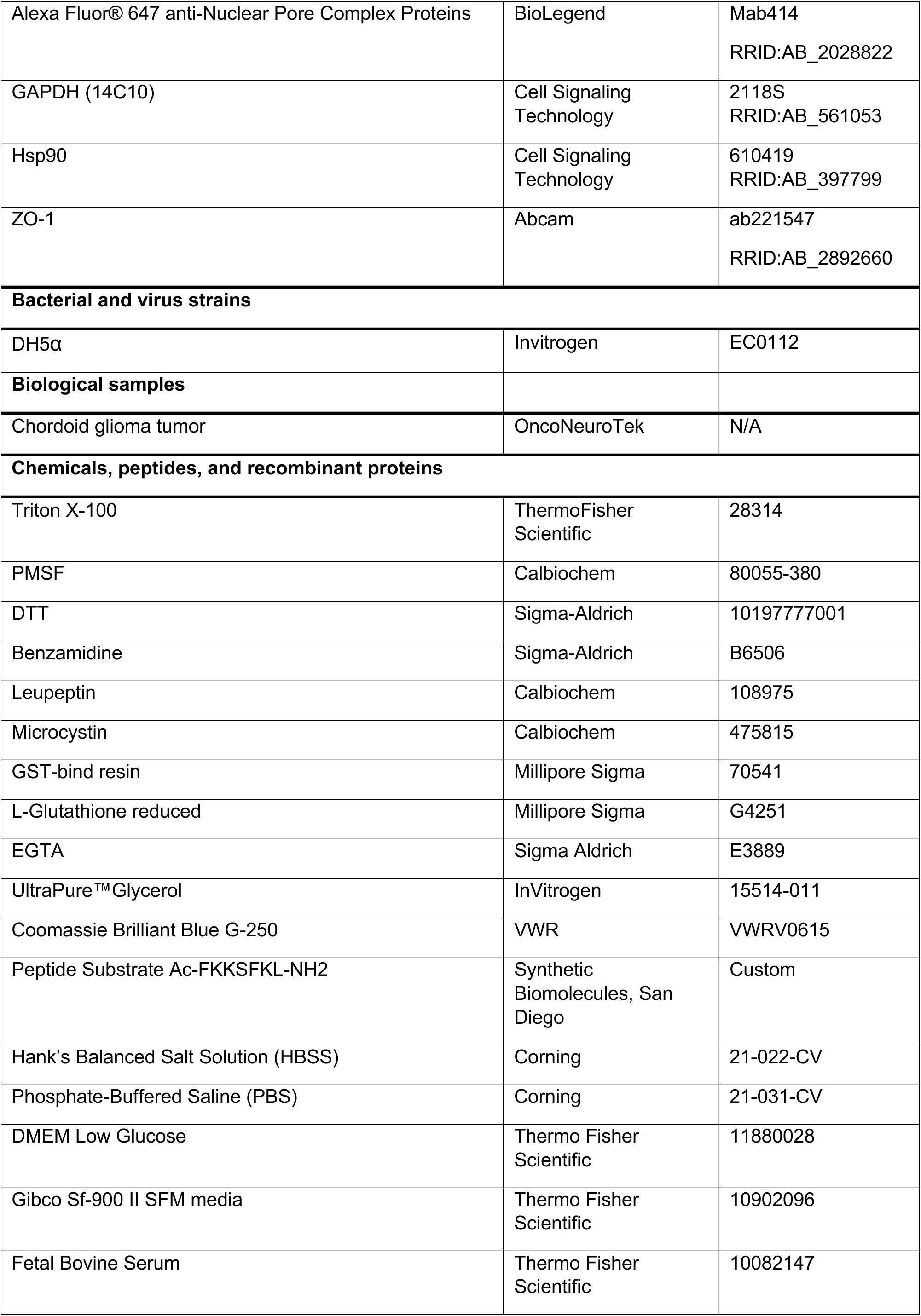
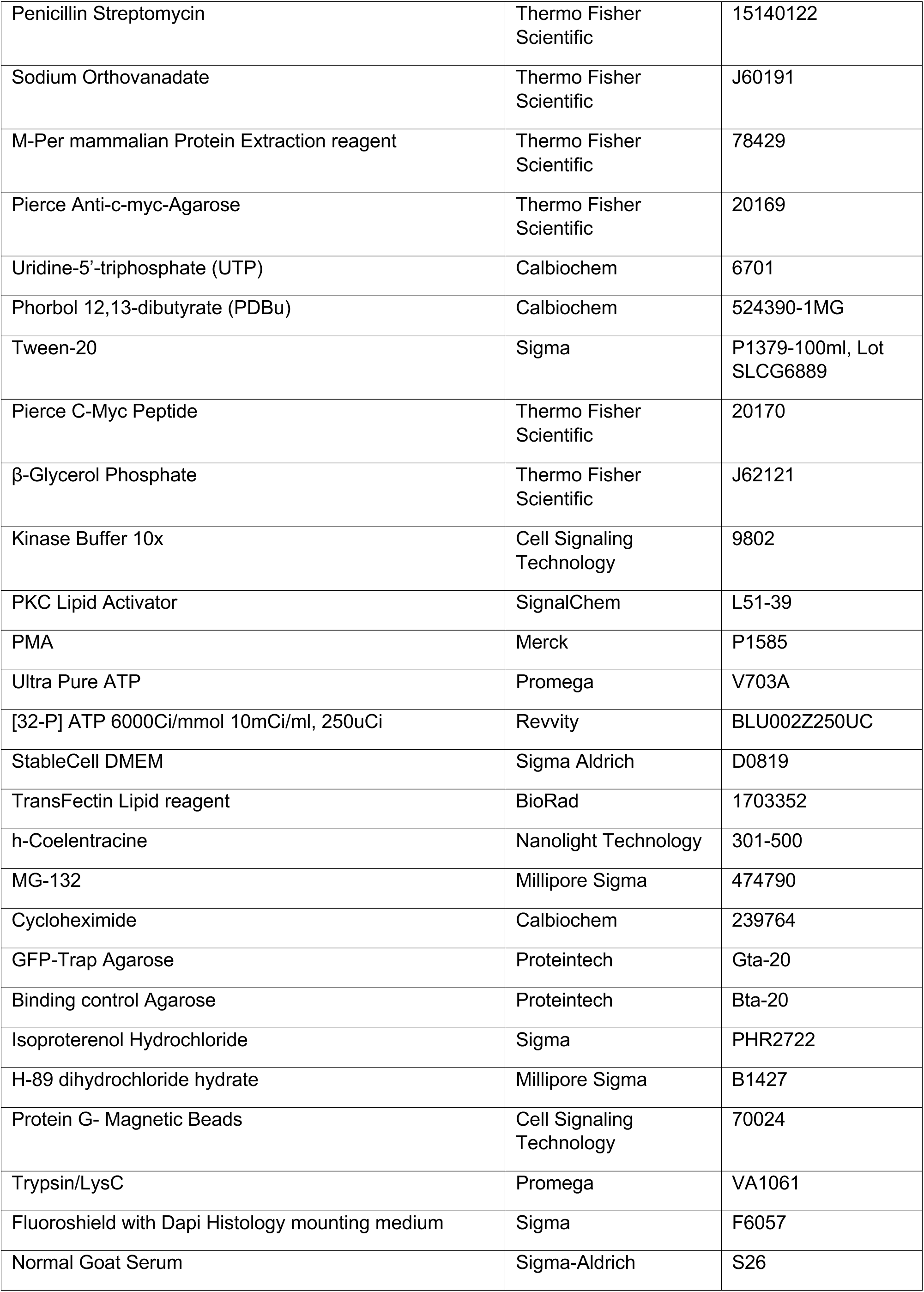
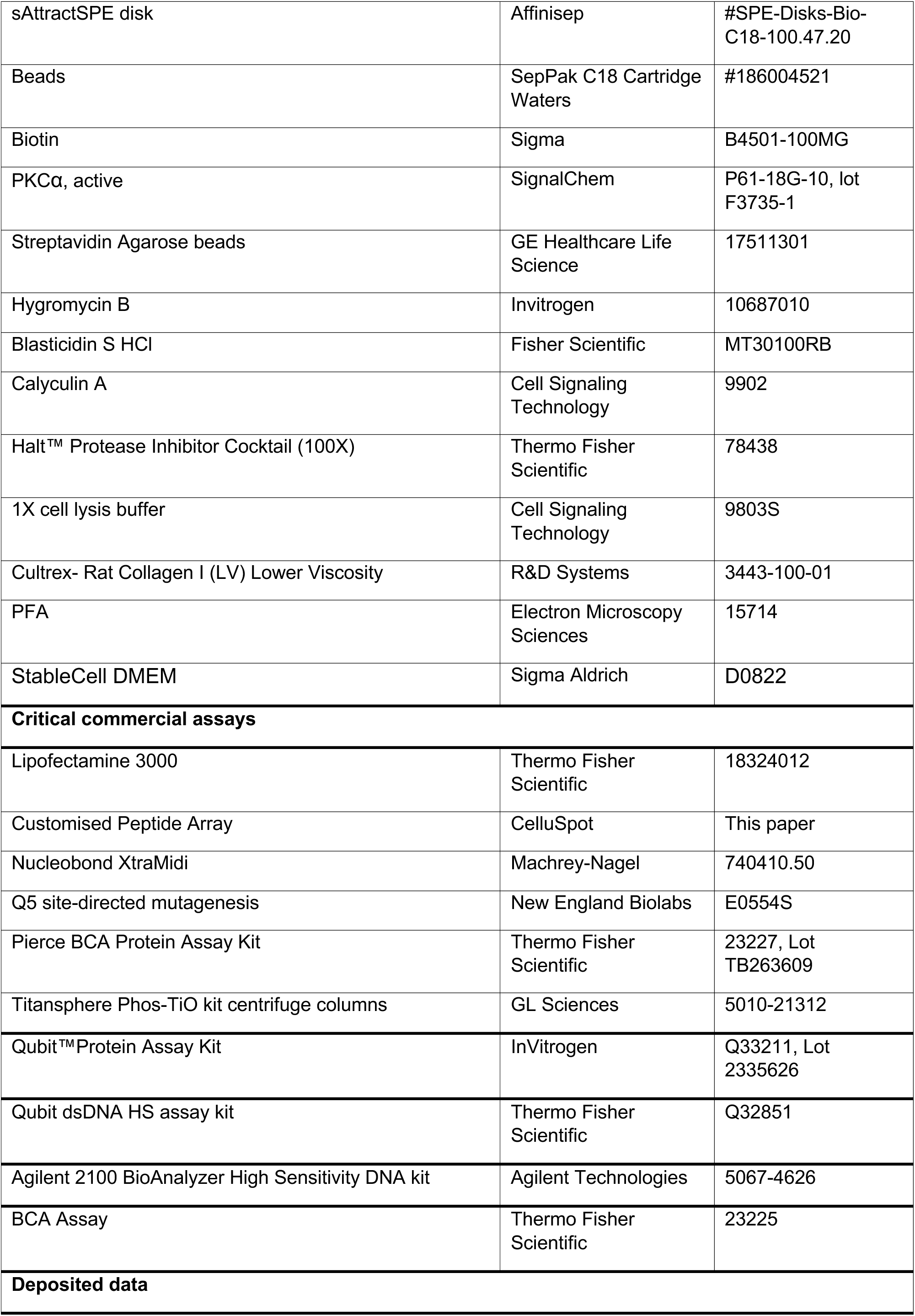
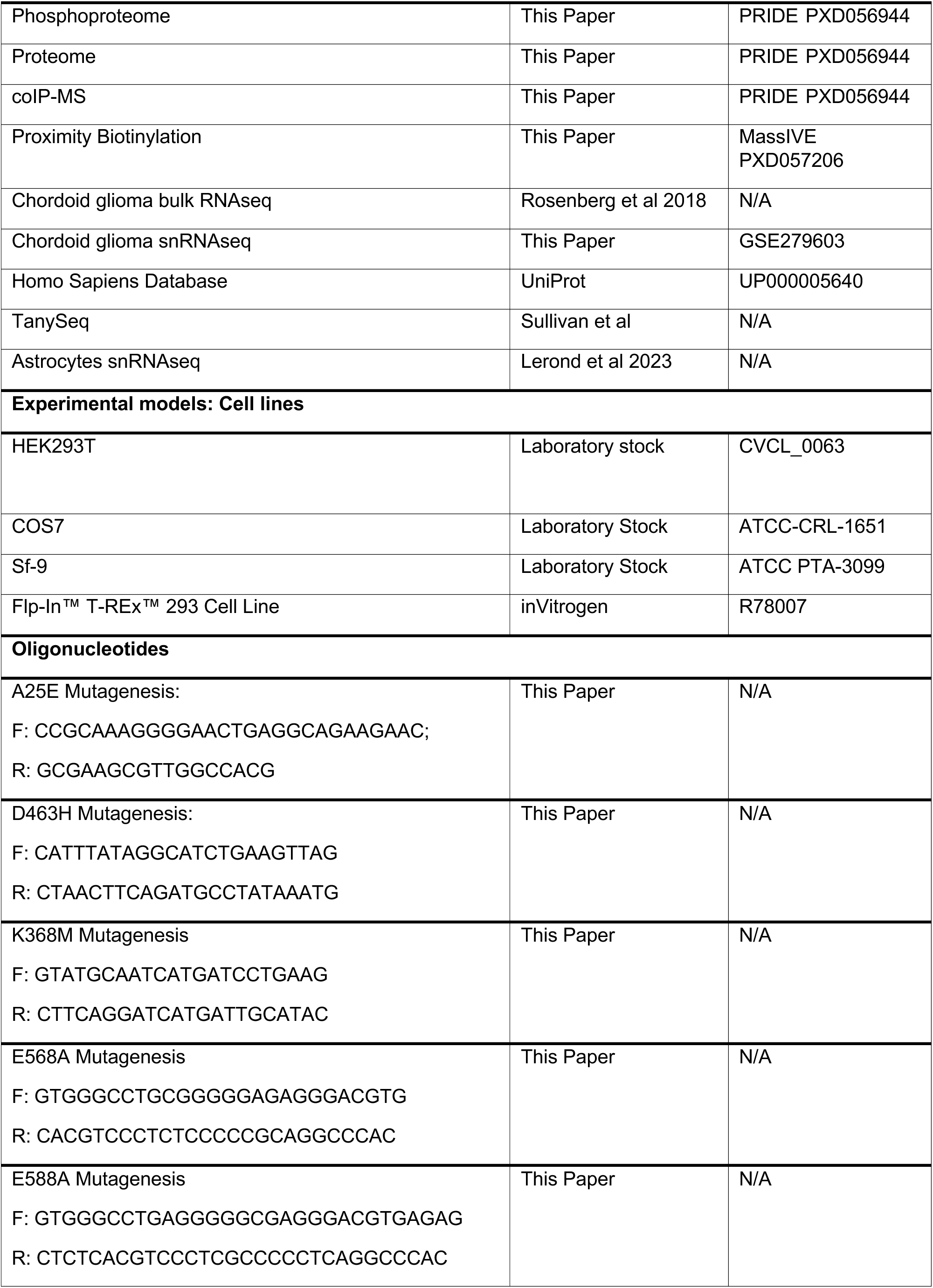
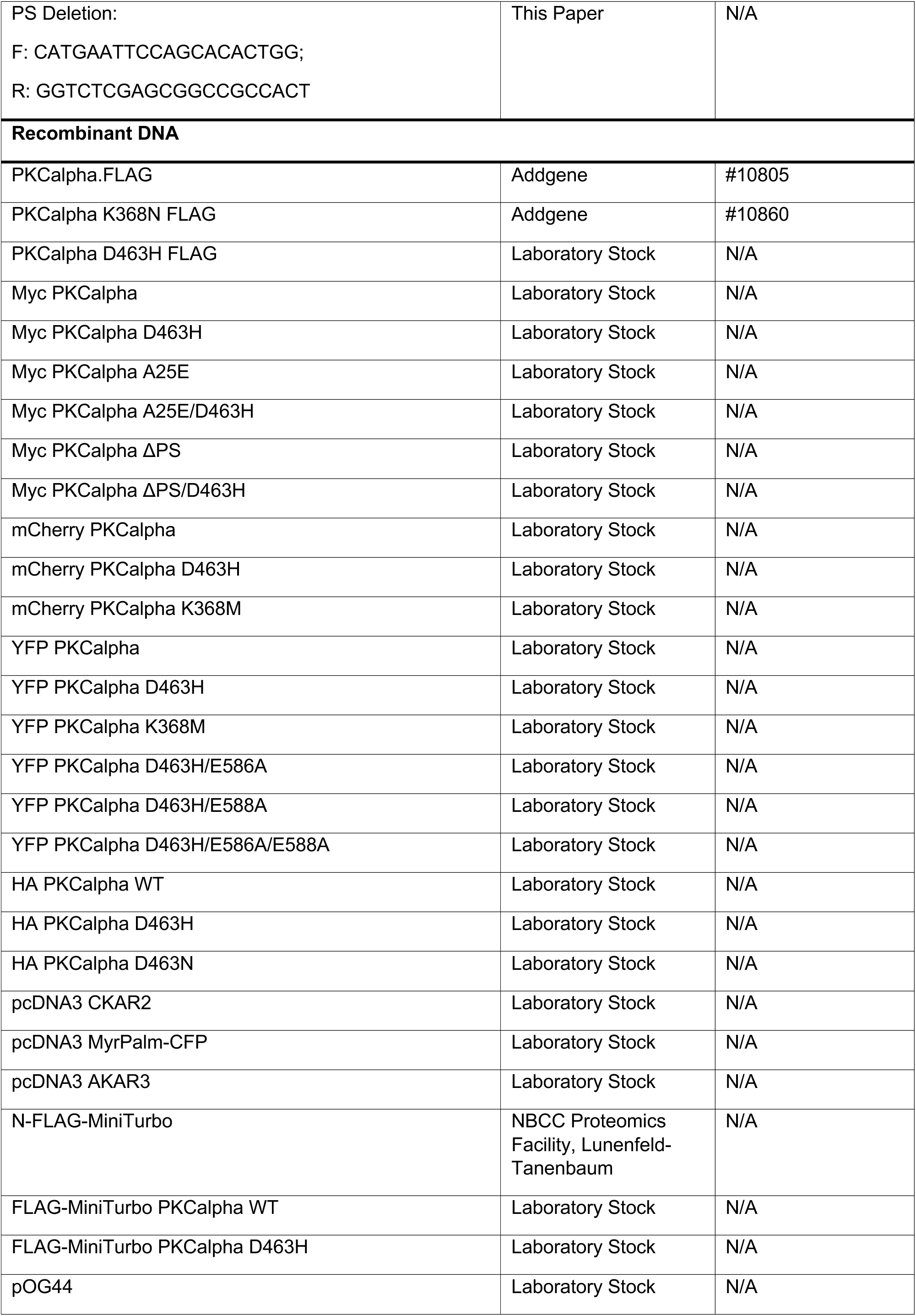
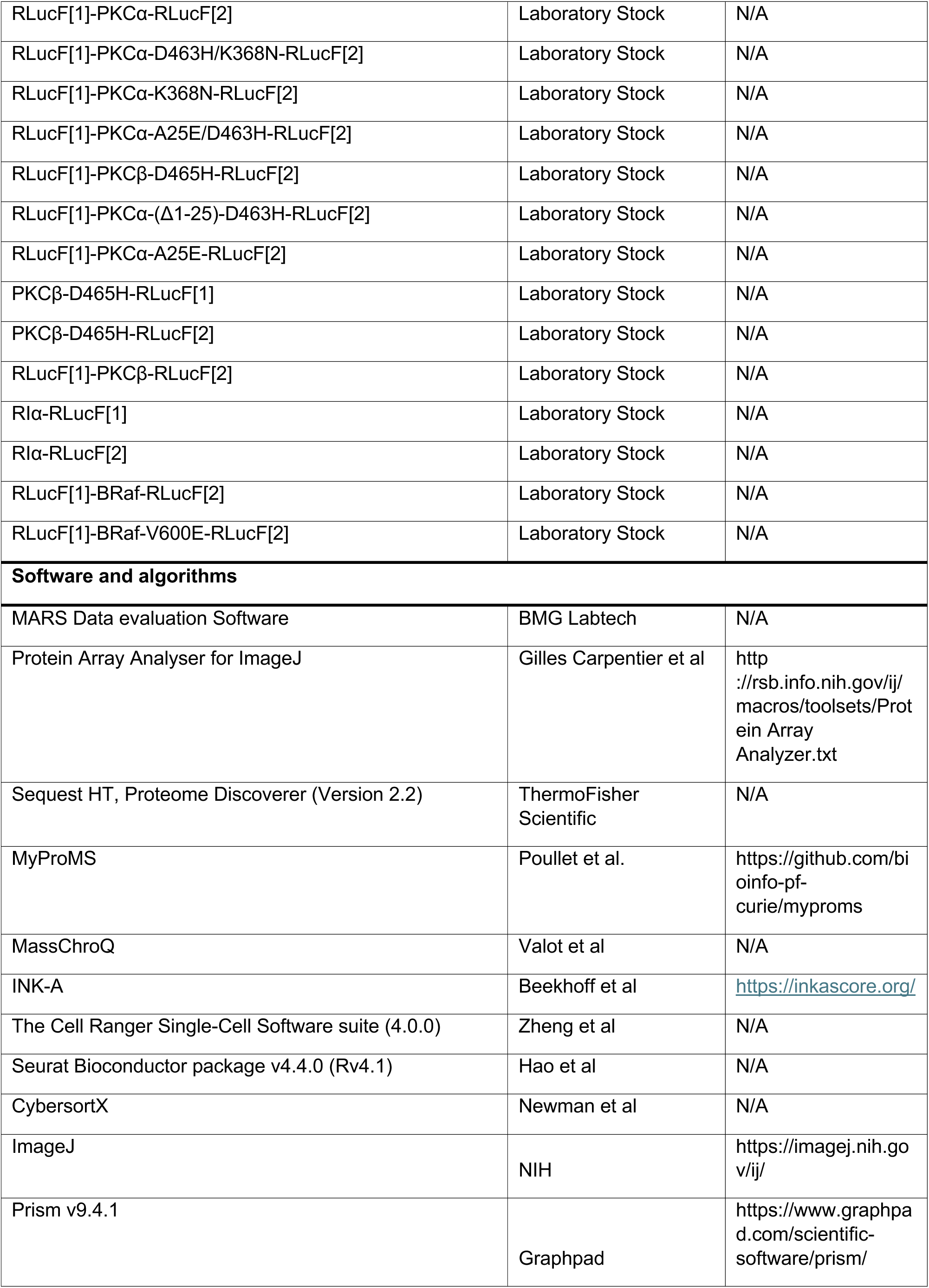
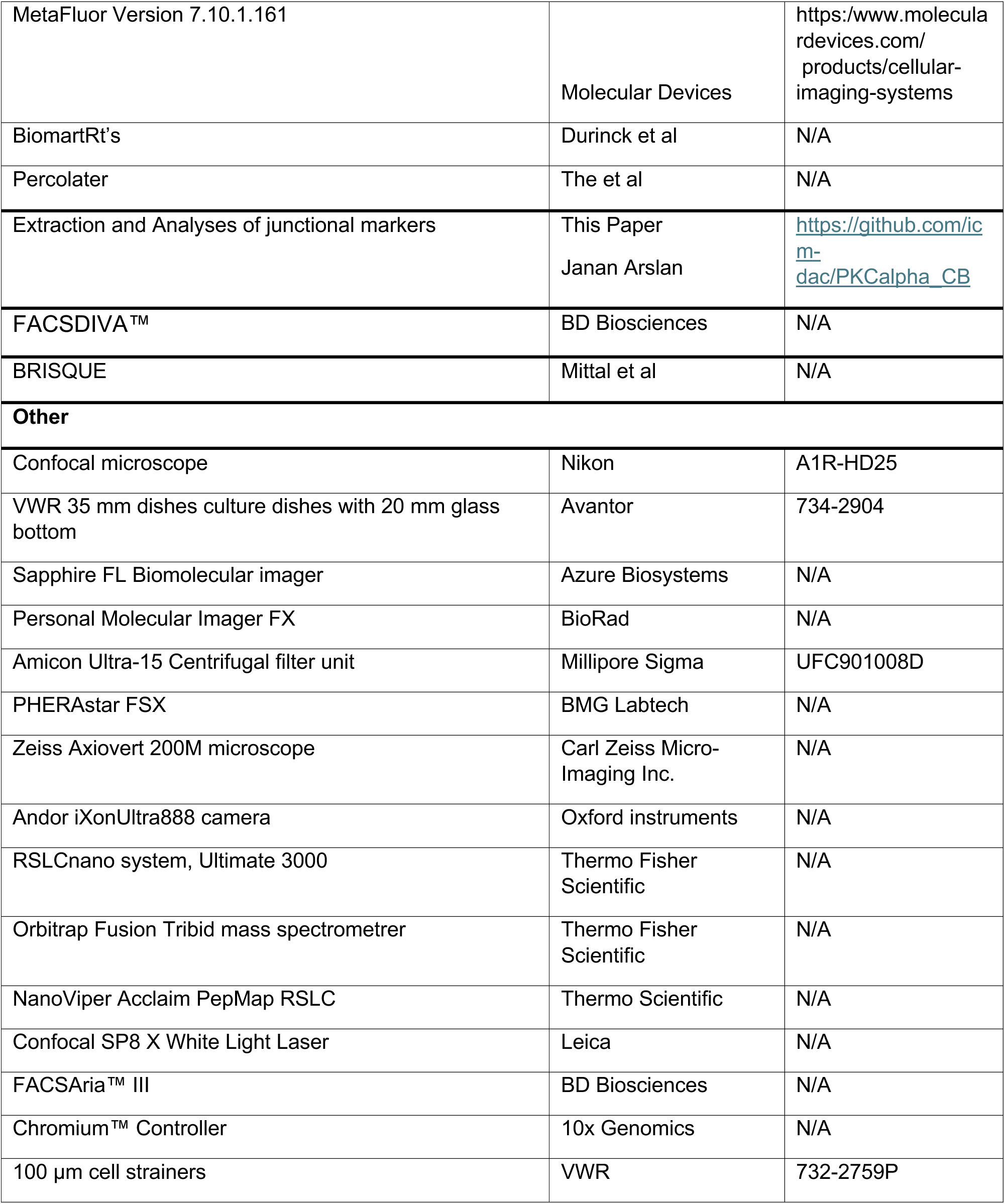

